# Task-dependent optimal representations for cerebellar learning

**DOI:** 10.1101/2022.08.15.504040

**Authors:** Marjorie Xie, Samuel Muscinelli, Kameron Decker Harris, Ashok Litwin-Kumar

## Abstract

The cerebellar granule cell layer has inspired numerous theoretical models of neural representations that support learned behaviors, beginning with the work of Marr and Albus. In these models, granule cells form a sparse, combinatorial encoding of diverse sensorimotor inputs. Such sparse representations are optimal for learning to discriminate random stimuli. However, recent observations of dense, low-dimensional activity across granule cells have called into question the role of sparse coding in these neurons. Here, we generalize theories of cerebellar learning to determine the optimal granule cell representation for tasks beyond random stimulus discrimination, including continuous input-output transformations as required for smooth motor control. We show that for such tasks, the optimal granule cell representation is substantially denser than predicted by classic theories. Our results provide a general theory of learning in cerebellum-like systems and suggest that optimal cerebellar representations are task-dependent.

## Introduction

A striking property of cerebellar anatomy is the vast expansion in number of granule cells compared to the mossy fibers that innervate them (Eccles et al., 1967). This anatomical feature has led to the proposal that the function of the granule cell layer is to produce a high-dimensional representation of lower-dimensional mossy fiber activity (Marr, 1969; Albus, 1971; Cayco-Gajic and Silver, 2019). In such theories of cerebellar cortex, granule cells integrate information from multiple mossy fibers and respond in a nonlinear manner to different input combinations. Detailed theoretical analysis has argued that anatomical parameters such as the ratio of granule cells to mossy fibers (Babadi and Sompolinsky, 2014), the number of inputs received by individual granule cells (Litwin-Kumar et al., 2017; Cayco-Gajic et al., 2017), and the distribution of granule-cell-to-Purkinje-cell synaptic weights (Brunel et al., 2004) have quantitative values that maximize the dimension of the granule cell representation and learning capacity. Sparse activity, which increases dimension, is also cited as a key property of the granule cell representation (Marr, 1969; Albus, 1971; Babadi and Sompolinsky, 2014; but see Spanne and Jörntell, 2015).

Theories that study the effects of dimension on learning typically focus on the ability of a system to perform categorization tasks with random, high-dimensional inputs (Barak et al., 2013; Babadi and Sompolinsky, 2014; Litwin-Kumar et al., 2017; Cayco-Gajic et al., 2017). In this case, increasing the dimension of the granule cell representation increases the number of inputs that can be discriminated (Rigotti et al., 2013; Litwin-Kumar et al., 2017). However, cerebellar circuits participate in diverse behaviors, including dexterous movements, inter-limb coordination, the formation of internal models, and cognitive behaviors (Ito and Itō, 1984; Wolpert et al., 1998; Strick et al., 2009). Cerebellum-like circuits, such as the insect mushroom body and the electrosensory system of electric fish, support other functions such as associative learning (Modi et al., 2020) and the cancellation of self-generated sensory signals (Kennedy et al., 2014), respectively. This diversity raises the question of whether learning high-dimensional categorization tasks is a sufficient framework for probing the function of granule cells and their analogs.

Indeed, several recent studies have reported dense activity in cerebellar granule cells in response to sensory stimulation or during motor control tasks (Knogler et al., 2017; Wagner et al., 2017; Giovannucci et al., 2017; Badura and De Zeeuw, 2017; Wagner et al., 2019), at odds with classic theories (Marr, 1969; Albus, 1971). Moreover, granule cell firing rates differ across cerebellar regions (Heath et al., 2014; Witter and De Zeeuw, 2015). In contrast to the dense activity in cerebellar granule cells, odor responses in Kenyon cells, the analogs of granule cells in the *Drosophila* mushroom body, are sparse, with 5–10% of neurons responding to odor stimulation (Turner et al., 2008; Honegger et al., 2011; Lin et al., 2014).

We propose that these differences can be explained by the capacity of representations with different levels of sparsity to support learning of different tasks. We show that, when learning input-output mappings for motor control tasks, the optimal granule cell representation is much denser than predicted by previous analyses. To explain this result, we develop a theory that predicts the performance of cerebellum-like circuits for arbitrary learning tasks. The theory describes how properties of cerebellar architecture and activity control these networks’ inductive bias: the tendency of a network toward learning particular types of input-output mappings (Sollich, 1998; Jacot et al., 2018; Bordelon et al., 2020; Canatar et al., 2021b; Simon et al., 2021). The theory shows that inductive bias, rather than the dimension of the representation alone, is necessary to explain learning performance across tasks. It also suggests that cerebellar regions specialized for different functions may adjust the sparsity of granule cell representations depending on the task. More broadly, we show that the sparsity of a neural code has a task-dependent influence on learning, a principle that may apply generally to sensorimotor representations across brain regions.

## Results

In our model, a granule cell layer of *M* neurons receives connections from a random subset of *N* mossy fiber inputs. Because *M* » *N* in the cerebellar cortex and cerebellum-like structures (approximately *M* = 200,000 and *N* = 7,000 for the neurons presynaptic to a single Purkinje cell in the cat brain; Eccles et al., 1967), we refer to the granule cell layer as the expansion layer and the mossy fiber layer as the input layer (Fig. 1A,B).

**Figure 1:**
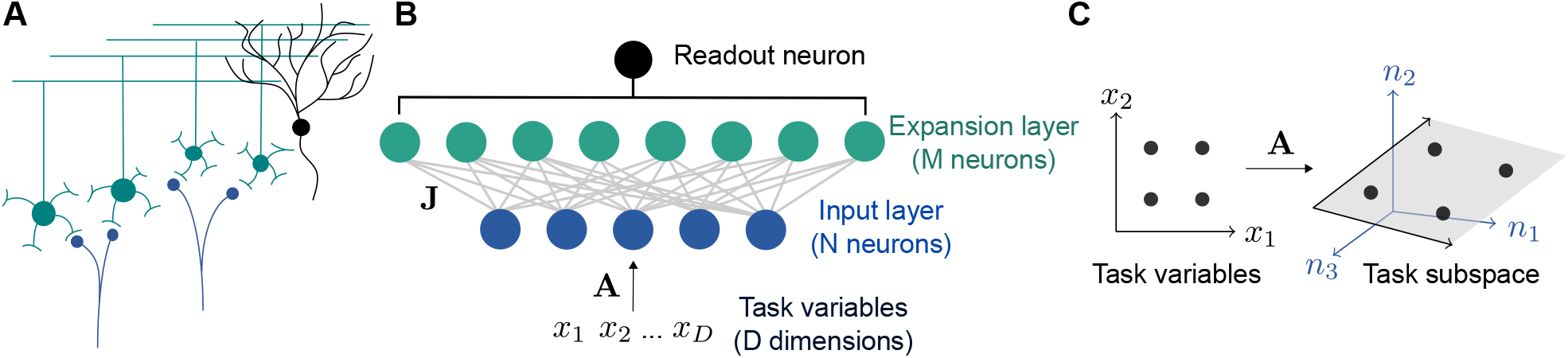
Schematic of cerebellar cortex model. A) Mossy fiber inputs (blue) project to granule cells (green), which send parallel fibers that contact a Purkinje cell (black). B) Diagram of neural network model. *D* task variables are embedded, via a matrix **A**, in the activity of *N* input layer neurons. Connections from the input layer to the expansion layer are described by a synaptic weight matrix **J**. C) Illustration of task subspace. Points **x** in a *D*-dimensional space of task variables are embedded in a *D*-dimensional subspace of the *N*-dimensional input layer activity **n** (*D*=2, *N* =3 illustrated).

A typical assumption in computational theories of the cerebellar cortex is that inputs are randomly distributed in a high-dimensional space (Marr, 1969; Albus, 1971; Brunel et al., 2004; Babadi and Sompolinsky, 2014; Billings et al., 2014; Litwin-Kumar et al., 2017). While this may be a reasonable simplification in some cases, many tasks, including cerebellum-dependent tasks, are likely best-described as being encoded by a low-dimensional set of variables. For example, the cerebellum is often hypothesized to learn a forward model for motor control (Wolpert et al., 1998), which uses sensory input and motor efference to predict an effector’s future state. Sources of motor efference copies include motor cortex, whose population activity lies on a low-dimensional manifold (Wagner et al., 2019; Huang et al., 2013; Churchland et al., 2010; Yu et al., 2009).

Therefore, we assume that the inputs to our model lie on a *D*-dimensional subspace embedded in the *N*-dimensional input space, where *D* is typically much smaller than *N* (Fig. 1B). We refer to this subspace as the “task subspace” (Fig. 1C). A location in this subspace is described by a *D* dimensional vector **x**, while the input layer activity is given by **n** = **Ax**, with **A** describing the embedding of the task variables in the input layer. An effective weight matrix **J**^eff^ = **JA**, which describes the selectivity of expansion layer neurons to task variables, is determined by **A** and the input-to-expansion-layer synaptic weight matrix **J**. The activity of neurons in the expansion layer is given by:

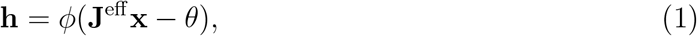

where *φ* is a threshold-linear activation function with threshold *θ*, and the nonlinearity and subtraction are applied element-wise. The threshold *θ* controls the coding level, which we denote by *f*, defined as the average fraction of neurons in the expansion layer that are active. We show results for *f* < 0.5, since extremely dense codes are rarely observed in experiments (Olshausen and Field, 2004; see Discussion). For analytical tractability, we begin with the case where the entries of **J**^eff^ are independent Gaussian random variables, as in previous theories (Rigotti et al., 2013; Barak et al., 2013; Babadi and Sompolinsky, 2014). Specifically, we assume that the columns of **A** are orthonormal (ensuring that the embedding of the task variables in the input layer preserves their geometry) and the entries of **J** are independent Gaussian random variables. Later, we will show that networks with more realistic connectivity exhibit similar behavior to this case.

### Optimal coding level is task-dependent

In our model, a learning task is defined by a mapping from task variables **x** to an output *f* (**x**), representing a target change in activity of a readout neuron, for example a Purkinje cell. The readout adjusts its incoming synaptic weights from the expansion layer to better approximate this target output. For example, for an associative learning task in which sensory stimuli are classified into categories such as appetitive or aversive, the task may be represented as a mapping from inputs to two discrete firing rates corresponding to the readout’s response to stimuli of each category (Fig. 2A). In contrast, for a forward model, in which the consequences of motor commands are computed using a model of movement dynamics, an input encoding the current sensorimotor state is mapped to a continuous output representing a predicted sensory variable (Fig. 2B).

**Figure 2:**
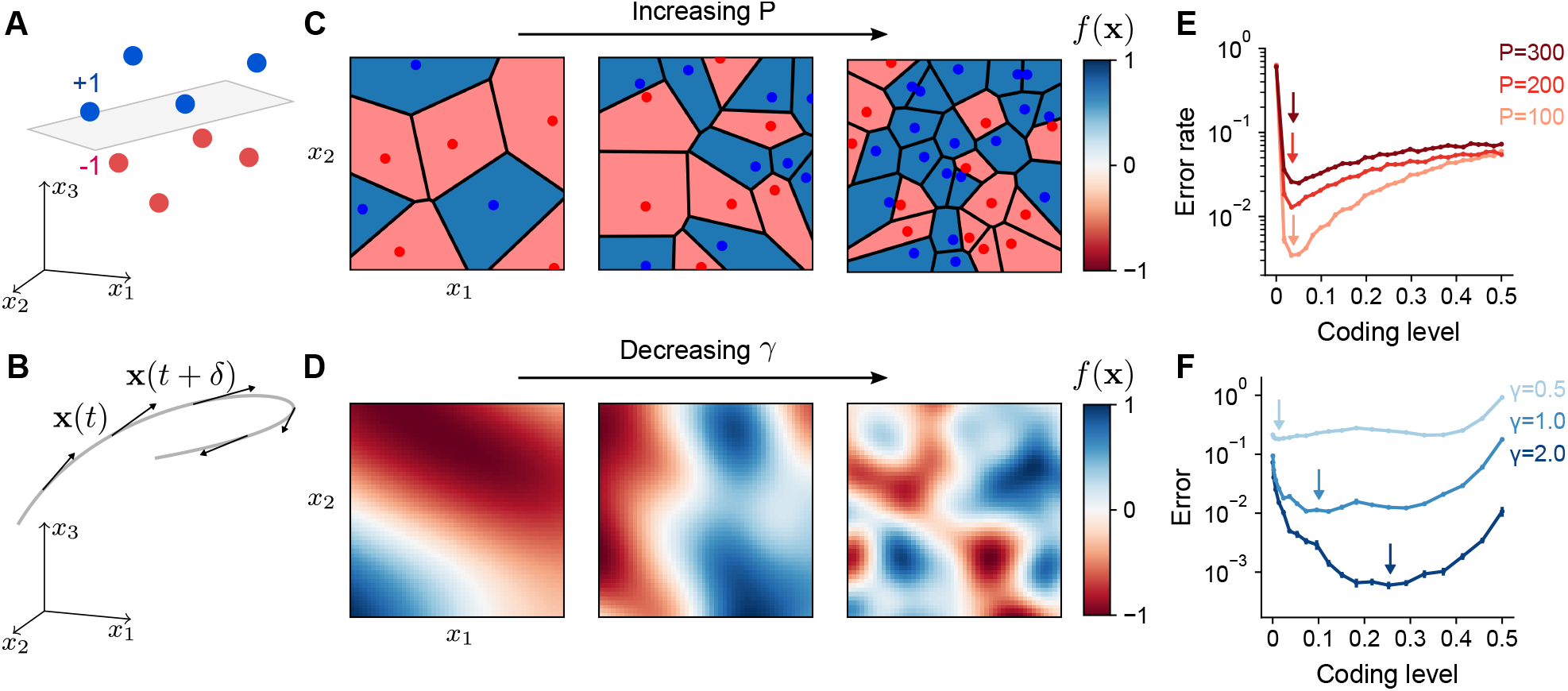
Optimal coding level depends on task. A) A random categorization task in which inputs are mapped to one of two categories (+1 or −1). Gray plane denotes the decision boundary of a linear classifier separating the two categories. B) A motor control task in which inputs are the sensorimotor states **x**(*t*) of an effector which change continuously along a trajectory (gray) and outputs are components of predicted future states *x*_*i*_(*t* + *δ*). C) Schematic of random categorization tasks with *P* input-category associations. The value of the target function *f* (**x**) (color) is a function of two task variables *x*_1_ and *x*_2_. D) Schematic of tasks involving learning a continuously varying Gaussian process target parameterized by a length scale *γ*. E) Error rate as a function of coding level for networks trained to perform random categorization tasks similar to (C). Arrows mark estimated locations of minima. F) Error as a function of coding level for networks trained to produce Gaussian process target functions of different length scales *γ* similar to (D). Standard error of the mean was computed across 20 realizations of network weights and tasks in (E) and 200 in (F).

To examine how properties of the expansion layer representation influence learning performance across tasks, we designed two families of tasks: one modeling categorization of random stimuli, which is often used to study the performance of expanded neural representations (Rigotti et al., 2013; Barak et al., 2013; Babadi and Sompolinsky, 2014; Litwin-Kumar et al., 2017; Cayco-Gajic et al., 2017), and the other modeling learning of a continuously varying output (Fig. 2C,D; see Methods). The former we refer to as a “random categorization task” and is parameterized by the number of input pattern-to-category associations *P*. The latter is parameterized by a length scale that determines how quickly the output changes as a function of the input (specifically, input-output functions are drawn from a Gaussian process with length scale *γ* for variations in *f* (**x**) as a function of **x**; see Methods). Later, we will also consider tasks implemented by specific cerebellum-like systems.

We trained the readout to approximate the target output and generalize to unseen inputs (see Methods). We began by examining the dependence of learning performance on the coding level of the expansion layer. For random categorization tasks, performance is maximized at low coding levels (Fig. 2E), consistent with previous results (Barak et al., 2013; Babadi and Sompolinsky, 2014). The optimal coding level remains below 0.1 in the model, regardless of the number of associations *P*, the level of input noise, and the dimension *D* (Supp. Fig. 1). For continuously varying outputs, the dependence is qualitatively different (Fig. 2F). The optimal coding level depends strongly on the length scale, with learning performance for slowly varying functions optimized at much higher coding levels than quickly varying functions. This dependence is at odds with previous theories of the role of sparse granule cell representations (Marr, 1969; Albus, 1971; Babadi and Sompolinsky, 2014; Billings et al., 2014) and shows that sparse activity does not always optimize performance for this broader set of tasks.

### Geometry of the expansion layer representation

To determine how the optimal coding level depends on the task, we begin by quantifying how the expansion layer transforms the geometry of the task subspace. Later we will address how this transformation affects the ability of the network to learn a target. For ease of analysis, we will assume for now that inputs are normalized, ∥**x**∥ = 1, so that they lie on the surface of a sphere in *D* dimensions. The set of neurons in the expansion layer activated by an input **x** are those neurons *i* for which the alignment of their effective weights with the input, 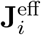. **x**, exceeds the activation threshold *θ* (Equation 1; Fig. 3A). Increasing *θ* reduces the size of this set of neurons and hence reduces the coding level.

**Figure 3:**
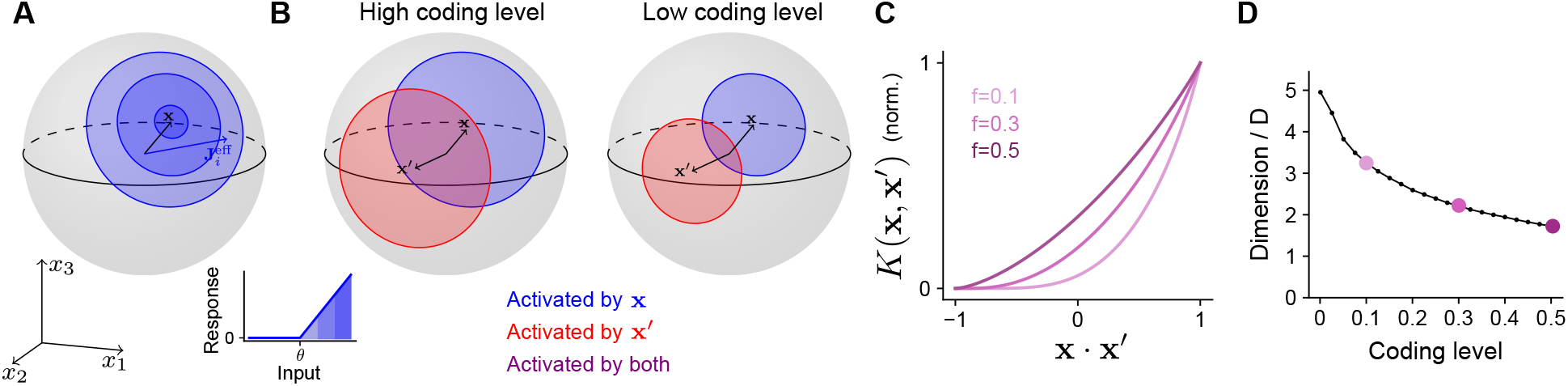
Effect of coding level on the expansion layer representation. A) Effect of activation threshold on coding level. A point on the surface of the sphere represents a neuron with effective weights 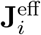. Blue region represents the set of neurons activated by **x**, i.e., neurons whose input exceeds the activation threshold *θ* (inset). Darker regions denote higher activation. B) Effect of coding level on the overlap between population responses to different inputs. Blue and red regions represent the neurons activated by **x** and **x**′, respectively. Overlap (purple) represents the set of neurons activated by both stimuli. High coding level leads to more active neurons and greater overlap. C) Kernel *K*(**x, x**′), normalized so that fully overlapping representations have an overlap of 1, plotted as a function of overlap in the space of task variables. The vertical axis corresponds to the ratio of the area of the purple region to the area of the red or blue regions in (B). D) Dimension of the expansion layer representation as a function of coding level for a finite-width network with *M* =10,000 and *D* = 3.

Different inputs activate different sets of neurons, and more similar inputs activate sets with greater overlap. As the coding level is reduced, this overlap is also reduced (Fig. 3B). In fact, this reduction in overlap is greater than the reduction in number of neurons that respond to either of the individual inputs, reflecting the fact that representations with low coding levels perform “pattern separation” (Fig. 3B, compare purple and red or blue regions).

This effect is summarized by the “kernel” of the network (Schölkopf and Smola, 2002; Rahimi and Recht, 2007), which measures overlap of representations in the expansion layer as a function of the task variables:

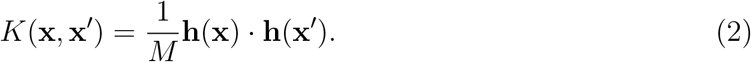

When inputs are normalized and the effective weights are Gaussian, we compute a semianalytic expression for the kernel of the expansion layer in the limit of a large expansion (*M* → ∞; see Appendix). In this case, the kernel depends only on the overlap of the task variables, *K*(**x, x**′) = *K*(**x** · **x**′). The resulting curves demonstrate that representations with lower coding levels exhibit greater pattern separation (Fig. 3C; Babadi and Sompolinsky, 2014). This is consistent with the observation that decreasing the coding level increases the dimension of the representation (Fig. 3D).

### Frequency decomposition of kernel and task explains optimal coding level

We now relate the geometry of the expansion layer representation to performance across the tasks we have considered. Previous studies focused on high-dimensional, random categorization tasks in which inputs belong to a small number of well-separated clusters and generalization to new inputs (“test patterns”) is assessed by adding noise to previously observed inputs (“training patterns”; Babadi and Sompolinsky, 2014; Litwin-Kumar et al., 2017; Fig. 4A). In this case, performance depends only on overlaps at two spatial scales: the overlap between training patterns belonging to different clusters, which is small, and the overlap between training and test patterns belonging to the same cluster, which is large (Fig. 4B). For such tasks, the kernel evaluated near these two values—specifically, the behavior of *K*(**x** · **x**′) near **x** · **x**′ = 0 and **x** · **x**′ = 1 − ∆, where ∆ is a measure of within-cluster noise—fully determines generalization performance (Fig. 4C; see Appendix). Sparse expansion layer representations reduce the overlap of patterns belonging to different clusters, increasing dimension and generalization performance (Fig. 3D).

**Figure 4:**
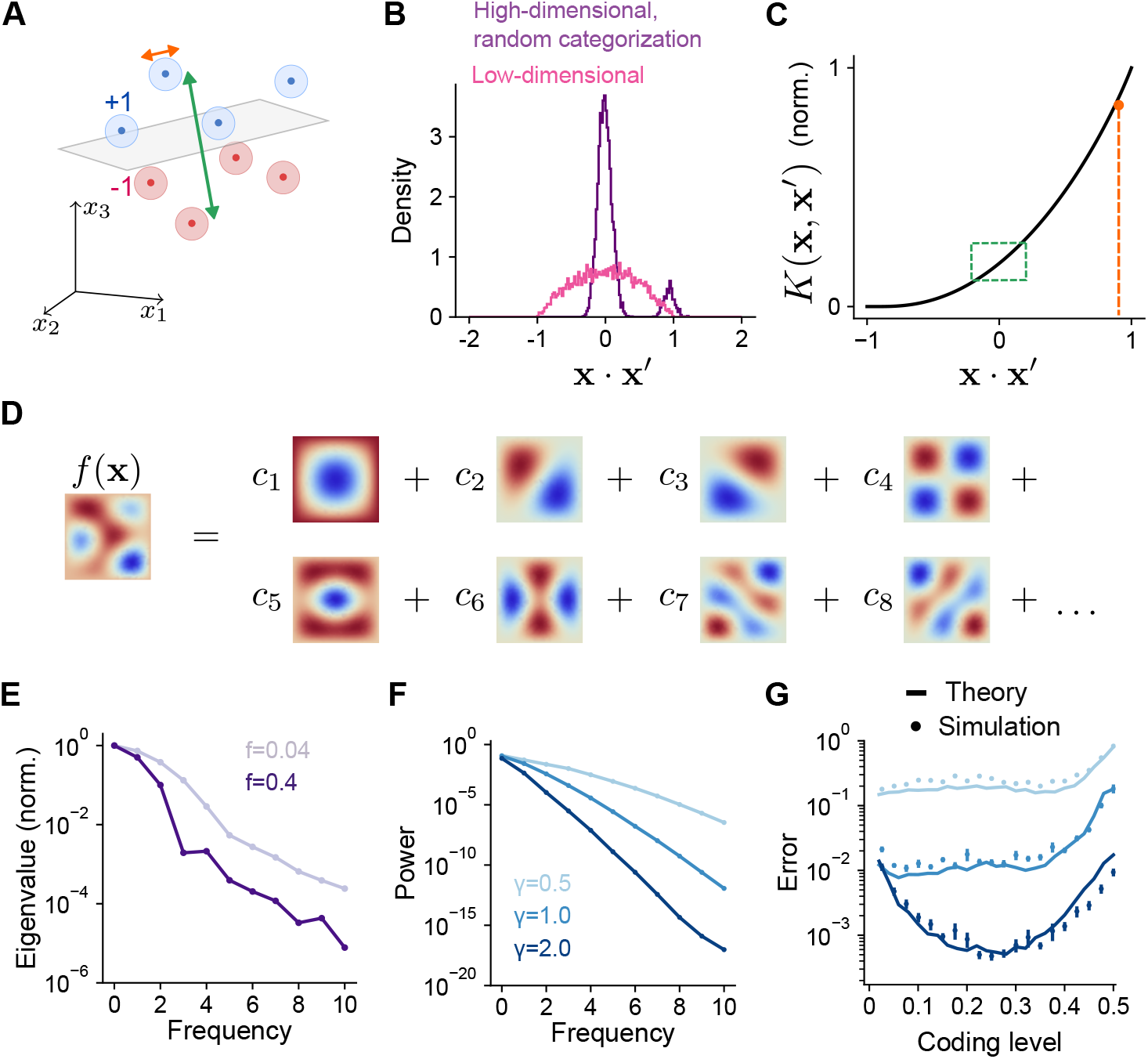
Frequency decomposition of network and target function. A) Geometry of high-dimensional categorization tasks where input patterns are drawn from random, noisy clusters (light regions). Performance depends on overlaps between training patterns from different clusters (green) and training and test patterns from the same cluster (orange). B) Distribution of overlaps of training and test patterns in the space of task variables for a high-dimensional task with random, clustered inputs as in (A) (purple) and a low-dimensional task with inputs drawn uniformly on a sphere in *D* dimensions (pink). C) Overlaps in (A) mapped onto the kernel function. Overlaps between training patterns from different clusters are small (green). Overlaps between training and test patterns from the same cluster are large (orange). D) Schematic illustration of basis function decomposition, for eigenfunctions on a square domain. E) Kernel eigenvalues (normalized by the sum of eigenvalues across modes) as a function of frequency for networks with different coding levels. F) Power as a function of frequency for Gaussian process target functions (same as Fig. 2F). Power is averaged over 20 realizations of target functions. G) Generalization error predicted using kernel eigenvalues (E) and target function decomposition (F) for the three tasks shown in (F). Standard error of the mean is computed across 10 realizations of network weights and tasks.

We study tasks where training patterns used for learning and test patterns used to assess generalization are both drawn according to a distribution over a low-dimensional space of task variables. Since overlaps between training and test patterns exhibit a broader distribution in this case (Fig. 4B), generalization performance depends on values of the kernel function evaluated across this entire range of overlaps. In particular, methods from the theory of kernel regression (Sollich, 1998; Jacot et al., 2018; Bordelon et al., 2020; Canatar et al., 2021b; Simon et al., 2021) quantify a network’s performance on a learning task by decomposing the target function into a set of basis functions (Fig. 4D). Performance is assessed by summing the contribution of each mode in this decomposition to generalization error.

The decomposition expresses the kernel as a sum of eigenfunctions weighted by eigenvalues, *K*(**x, x**′) = ∑_*α*_ *λ*_*α*_*ψ*_*α*_(**x**)*ψ*_*α*_(**x**′). The eigenfunctions are determined by the network architecture and the distribution of inputs. As we show below, the eigenvalues *λ*_*α*_ determine the ease with which each corresponding eigenfunction *ψ*_*α*_(**x**)—one element of the basis function decomposition—is learned by the network. Under our present assumptions of Gaussian effective weights and uniformly distributed, normalized input patterns, the eigenfunctions are the spherical harmonic functions. These functions are ordered by increasing frequency, with higher frequencies corresponding to functions that vary more quickly as a function of the task variables. Spherical harmonics are defined for any input dimension; for example, in two dimensions they are the Fourier modes. We find that coding level substantially changes the frequency dependence of the eigenvalues associated with these eigenfunctions (Fig. 4E). Higher coding levels increase the relative magnitude of the low frequency eigenvalues compared to high frequency eigenvalues. As we will show, this results in a different inductive bias for networks with different coding levels.

To calculate learning performance for an arbitrary task, we decompose the target function in the same basis as that of the kernel:

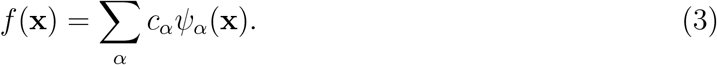

The coefficient *c*_*α*_ quantifies the weight of mode *α* in the decomposition. For the Gaussian process targets we have considered, increasing length scale corresponds to a greater relative contribution of low versus high frequency modes (Fig. 4F). Using these coefficients and the eigenvalues (Fig. 4E), we obtain an analytical prediction of the mean-squared generalization error (“error”) for learning any given task (Fig. 4G; see Methods):

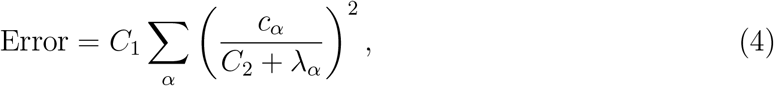

where *C*_1_ and *C*_2_ do not depend on *α* (Canatar et al., 2021b; Simon et al., 2021; see Methods). Equation (4) illustrates that for equal values of *c*_*α*_, modes with greater *λ*_*α*_ contribute less to the generalization error.

Our theory reveals that the optima observed in Fig. 2F are a consequence of the difference in eigenvalues of networks with different coding levels. This reflects an inductive bias (Sollich, 1998; Jacot et al., 2018; Bordelon et al., 2020; Canatar et al., 2021b; Simon et al., 2021) of networks with low and high coding levels toward the learning of high and low frequency functions, respectively (Fig. 4E, Supp. Fig. 2). Thus, the coding level’s effect on a network’s inductive bias, rather than dimension alone, determines learning performance. Previous studies that focused only on random categorization tasks did not observe this dependence, since errors in such tasks are dominated by the learning of high frequency components, for which sparse activity is optimal (Supp. Fig. 3).

### Performance of sparsely connected expansions

To simplify our analysis, we have so far assumed full connectivity between input and expansion layers without a constraint on excitatory or inhibitory synaptic weights. In particular, we have assumed that the effective weight matrix **J**^eff^ contains independent Gaussian entries (Fig. 5A, top). However, synaptic connections between mossy fibers and granule cells are sparse and excitatory (Sargent et al., 2005), with a typical in-degree of *K* = 4 mossy fibers per granule cell (Fig. 5A, bottom). We therefore analyzed the performance of model networks with more realistic connectivity. Surprisingly, when **J** is sparse and nonnegative, both overall generalization performance and the task-dependence of optimal coding level remain unchanged (Fig. 5B).

**Figure 5:**
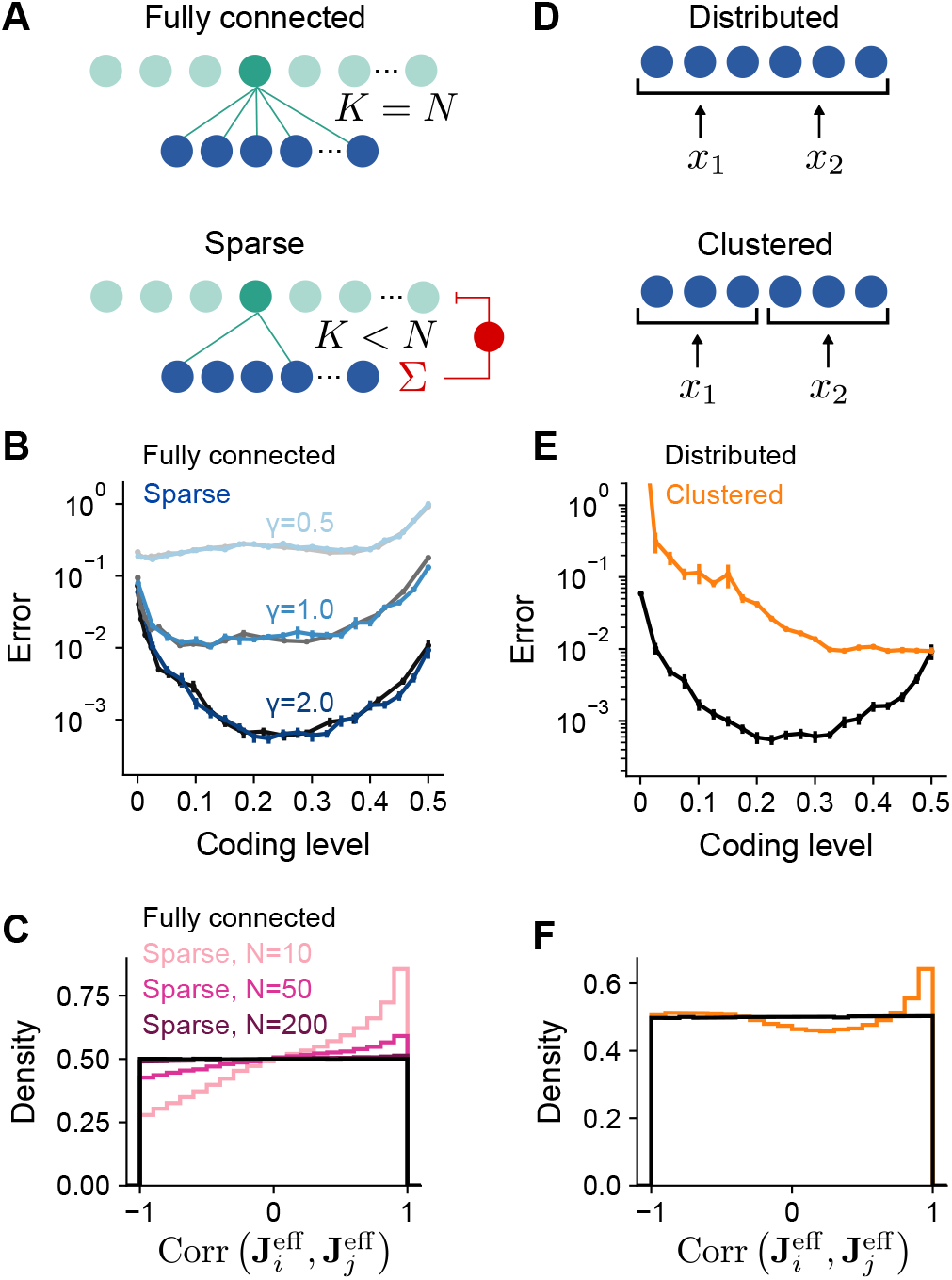
Performance of networks with sparse connectivity. A) Top: Fully connected network. Bottom: Sparsely connected network with in-degree *K* < *N* and excitatory weights with global inhibition onto expansion layer neurons. B) Error as a function of coding level for fully connected Gaussian weights (black) and sparse excitatory weights (blue). Target functions are Gaussian processes as in Fig. 2F. C) Distributions of synaptic weight correlations Corr 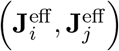, where 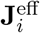 is the *i*th row of **J**^eff^, for pairs of expansion layer neurons in networks with different numbers of input layer neurons *N* (colors) when *K* = 4 and *D* = 3. Black distribution corresponds to fully connected network with Gaussian weights. D) Schematic of the selectivity of input layer neurons to task variables in distributed and clustered representations. E) Error as a function of coding level for networks with distributed (black, same as in B) and clustered (orange) representations. F) Distributions of Corr 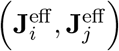 for pairs of expansion layer neurons in networks with distributed and clustered input representations when *K* = 4, *D* = 3, and *N* = 1000. Standard error of the mean was computed across 200 realizations of network weights and tasks in (B) and 100 in (E), orange curve.

To understand this result, we examined how **J** and **A** shape the statistics of the effective weights onto the expansion layer neurons **J**^eff^. A desirable property of the expansion layer representation is that these effective weights sample the space of task variables uniformly (Fig. 3A), increasing the heterogeneity of tuning of expansion layer neurons (Litwin-Kumar et al., 2017). This occurs when **J**^eff^ is a matrix of independent random Gaussian entries. In particular, if the columns of **A** are orthornormal and **J** is fully-connected with independent Gaussian entries, **J**^eff^ indeed has this property.

However, when **J** is sparse and nonnegative, expansion layer neurons that share connections from the same input layer neurons receive correlated input currents. When *N* is small and **A** is random, fluctuations in **A** lead to biases in the input layer’s sampling of task variables which are inherited by the expansion layer. We quantify this by computing the distribution of correlations between the effective weights for pairs of expansion layer neurons, Corr 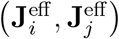. This distribution indeed deviates from uniform sampling when *N* is small (Fig. 5C). However, even when *N* is moderately large (but much less than *M*), only small deviations from uniform sampling of task variables occur for low dimensional tasks as long as *D* < *K* ≪ *N* (see Appendix). In contrast, for high-dimensional tasks (*D* ∼ *N*), *K* ≪ *D* is sufficient, in agreement with previous findings (Litwin-Kumar et al., 2017). For realistic cerebellar parameters (*N* = 7,000 and *K* = 4), the distribution is almost indistinguishable from that corresponding to uniform sampling (Fig. 5C), consistent with the similar learning performance of these two cases (Fig. 5B).

In the above analysis, an important assumption is that **A** is dense and random, so that the input layer forms a distributed representation in which each input layer neuron responds to a random combination of task variables (Fig. 5D, top). If, on the other hand, the input layer forms a clustered representation containing groups of neurons that each encode a single task variable (Fig. 5D, bottom), we may expect different results. Indeed, with a clustered representation, sparse connectivity dramatically reduces performance (Fig. 5E). This is because the distribution of Corr 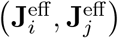 deviates substantially from that corresponding to uniform sampling (Fig. 5F), even as *N* → ∞ (see Appendix). Specifically, increasing *N* does not reduce the probability of two expansion layer neurons being connected to input layer neurons that encode the same task variables and therefore receiving highly correlated currents. As a result, expansion layer neurons do not sample task variables uniformly and performance is dramatically reduced.

Our results show that networks with small *K*, moderately large *N*, and a distributed input layer representation approach the performance of networks that sample task variables uniformly. This equivalence validates the applicability of our theory to these more realistic networks. It also argues for the importance of distributed sensorimotor representations in the cortico-cerebellar pathway, consistent with the distributed nature of representations in motor cortex (Shenoy et al., 2013; Muscinelli et al., 2022).

### Optimal cerebellar architectures are consistent across tasks

A history of theoretical modeling has shown a remarkable correspondence between anatomical properties of the cerebellar cortex and model parameters optimal for learning. These include the in-degree *K* of granule cells (Marr, 1969; Litwin-Kumar et al., 2017; Cayco-Gajic et al., 2017), the expansion ratio of the granule cells to the mossy fibers *M/N* (Babadi and Sompolinsky, 2014; Litwin-Kumar et al., 2017), and the distribution of synaptic weights from granule cells to Purkinje cells (Brunel et al., 2004; Clopath et al., 2012; Clopath and Brunel, 2013). In these studies, model performance was assessed using random categorization tasks. We have shown that optimal coding level is dependent on the task being learned, raising the question of whether optimal values of these architectural parameters are also task-dependent.

Sparse connectivity (*K*=4, consistent with the typical in-degree of cerebellar granule cells) has been shown to optimize learning performance in cerebellar cortex models (Litwin-Kumar et al., 2017; Cayco-Gajic et al., 2017). We examined the performance of networks with different granule cell in-degrees learning Gaussian process targets. The optimal in-degree is small for all the tasks we consider, suggesting that sparse connectivity is sufficient for high performance across a range of tasks (Fig. 6A). This is consistent with the previous observation that the performance of a sparsely connected network approaches that of a fully connected network (Fig. 5B).

**Figure 6:**
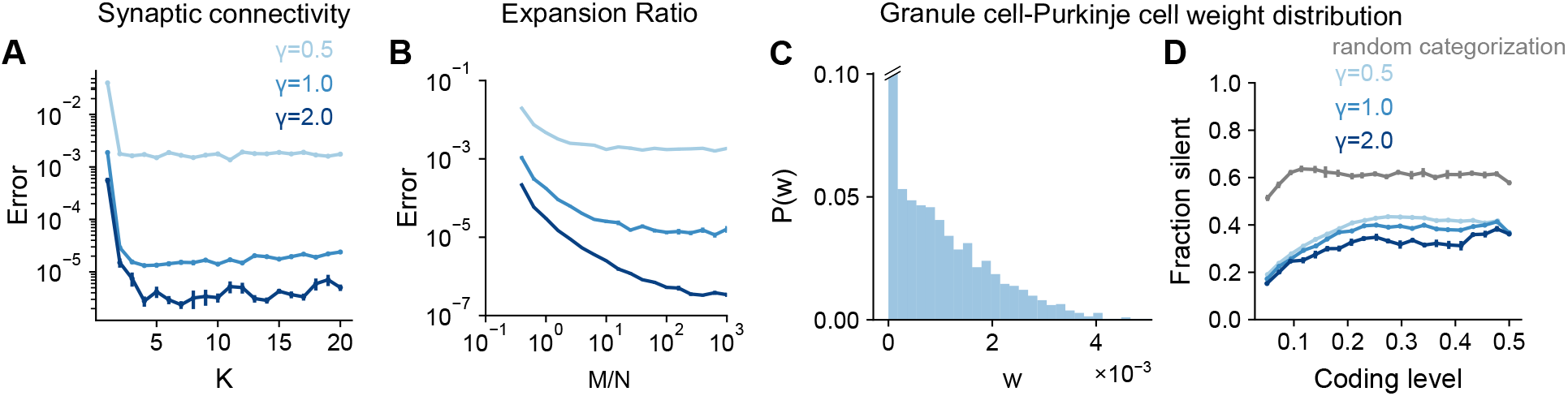
Task-independence of optimal anatomical parameters. A) Error as a function of in-degree *K* for networks learning Gaussian process targets. The total number of synaptic connections *S* = *MK* is held constant. This constraint introduces a trade-off between having many neurons with small synaptic degree or fewer neurons with large synaptic degree (Litwin-Kumar et al., 2017). *S* = 10^4^, *D* = 3, *f* = 0.3. B) Error as a function of expansion ratio *M/N* for networks learning Gaussian process targets. *D* = 3, *N* = 700, *f* = 0.3. C) Distribution of granule-cell-to-Purkinje cell weights for a network trained on nonnegative Gaussian process targets with *f* = 0.3, *D* = 3, *γ* = 1. Granule-cell-to-Purkinje cell weights are constrained to be nonnegative (Brunel et al., 2004). D) Fraction of granule-cell-to-Purkinje cell weights that are silent in networks learning nonnegative Gaussian process targets (blue) and random categorization tasks (gray).

Previous studies also showed that the expansion ratio from mossy fibers to granule cells *M/N* controls the dimension of the granule cell representation (Babadi and Sompolinsky, 2014; Litwin-Kumar et al., 2017). The dimension increases with expansion ratio but saturates as expansion ratio approaches the anatomical value (*M/N* ≈ 30 for the inputs presynaptic to an individual Purkinje cell when *f* ≈ 0.1). These studies assumed that mossy fiber activity is uncorrelated (*D* = *N*) rather than low-dimensional (*D* < *N*). This raises the question of whether a large expansion is beneficial when *D* is small. We find that when the number of training patterns *P* is sufficiently large, performance still improves as *M/N* approaches its anatomical value, showing that Purkinje cells can exploit their large number of presynaptic inputs even in the case of low-dimensional inputs (Fig. 6B).

Brunel et al. (2004) showed that the distribution of granule-to-Purkinje cell synaptic weights is consistent with the distribution that maximizes the number of random binary input-output mappings stored. This distribution exhibits a substantial fraction of silent synapses, consistent with experiments. These results also hold for analog inputs and outputs (Clopath and Brunel, 2013) and for certain forms of correlations among binary inputs and outputs (Clopath et al., 2012). However, the case we consider, where targets are a smoothly varying function of task variables, has not been explored. We observe a similar weight distribution for these tasks (Fig. 6C), with the fraction of silent synapses remaining high across coding levels (Fig. 6D). The fraction of silent synapses is lower for networks learning Gaussian process targets than those learning random categorization tasks, consistent with the capacity of a given network for learning such targets being larger (Clopath et al., 2012).

Although optimal coding level is task-dependent, these analyses suggest that optimal architectural parameters are largely task-independent. Thus, whereas coding level tunes the inductive bias of the network to favor the learning of specific tasks, these architectural parameters control properties of the representation that improve performance across tasks. In particular, sparse in-degree and a large expansion support uniform sampling of low-dimensional task variables (consistent with Fig. 5C), while a large fraction of silent synapses is a consequence of a readout that maximizes learning performance (Brunel et al., 2004).

### Modeling specific behaviors dependent on cerebellum-like structures

So far, we have considered analytically tractable families of tasks with parameterized input-output functions. Next, we extend our results to realistic tasks constrained by the functions of specific cerebellum-like systems, which include both highly structured, continuous input-output mappings and random categorization tasks.

To model the cerebellum’s role in predicting the consequences of motor commands (Wolpert et al., 1998), we examined the optimal coding level for learning the dynamics of a two-joint arm (Fagg et al., 1997). Given an initial state, the network predicts the change in the future position of the arm (Fig. 7A). Performance is optimized at substantially higher coding levels than for random categorization tasks, consistent with our previous results for continuous input-output mappings (Fig. 2E,F).

**Figure 7:**
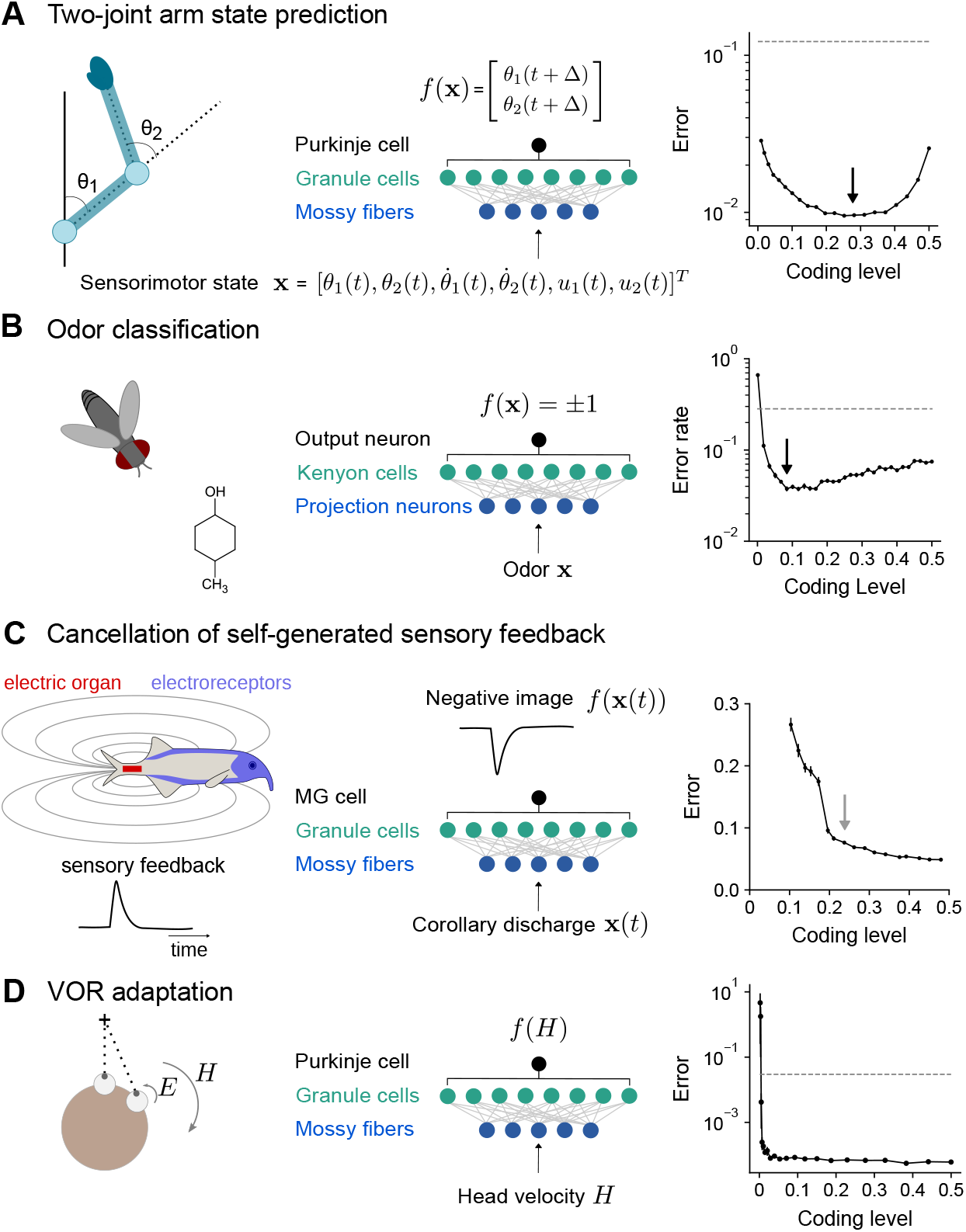
Optimal coding level across tasks and neural systems. A) Left: Schematic of two-joint arm. Center: Cerebellar cortex model in which sensorimotor task variables at time *t* are used to predict hand position at time *t* + *δ*. Right: Error as a function of coding level. Black arrow indicates location of optimum. Dashed line indicates performance of a readout of the input layer. B) Left: Odor categorization task. Center: *Drosophila* mushroom body model in which odors activate olfactory projection neurons and are associated with a binary category (appetitive or aversive). Right: Error rate, similar to (A), right. C) Left: Schematic of electrosensory system of the mormyrid electric fish, which learns a negative image to cancel the self-generated feedback from electric organ discharges sensed by electroreceptors. Center: Electrosensory lateral line lobe (ELL) model in which MG cells learn a negative image. Right: Error as a function of coding level. Gray arrow indicates location of coding level estimated from biophysical parameters (Kennedy et al., 2014). D) Left: Schematic of the vestibulo-cular reflex (VOR). Head rotations with velocity *H* trigger eye motion in the opposite direction with velocity *E*. During VOR adaptation, organisms adapt to different gains (*E/H*). Center: Cerebellar cortex model in which target function is Purkinje cell tuning to head velocity. Right: Error, similar to (A), right.

The mushroom body, a cerebellum-like structure in insects, is required for learning of associations between odors and appetitive or aversive valence (Modi et al., 2020). This behavior can be represented as a mapping from random representations of odors in the input layer to binary category labels (Fig. 7B). The optimal coding level in a model with parameters consistent with the *Drosophila* mushroom body is less than 0.1, consistent with our previous results for random categorization tasks (Fig. 2E) and the sparse odor-evoked responses in *Drosophila* Kenyon cells (Turner et al., 2008; Honegger et al., 2011; Lin et al., 2014).

The prediction and cancellation of self-generated sensory feedback has been studied extensively in mormyrid weakly electric fish and depends on the electrosensory lateral line lobe (ELL), a cerebellum-like structure (Bell et al., 2008). Granule cells in the ELL provide a temporal basis for generating negative images that are used to cancel self-generated feedback (Fig. 7C). We extended a detailed model of granule cells and their inputs (Kennedy et al., 2014) to study the influence of coding level on the effectiveness of this basis. The performance of this model saturated at relatively high coding levels, and notably the coding level corresponding to biophysical parameters estimated from data coincided with the value at which further increases in performance were modest. This observation suggests that coding level is also optimized for task performance in this system.

A canonical function of the mammalian cerebellum is the adjustment of the vestibulo-ocular reflex (VOR), in which motion of the head is detected and triggers compensatory ocular motion in the opposite direction. During VOR learning, Purkinje cells are tuned to head velocity, and their tuning curves are described as piecewise linear functions (Lisberger et al., 1994) (Fig. 7D). Although *in vivo* population recordings of granule cells during VOR adaptation are not, to our knowledge, available for comparison, our model predicts that performance for learning such tuning curves is high across a range of coding levels and shows that sparse codes are sufficient (although not necessary) for such tasks (Fig. 7A).

These results predict diverse coding levels across different behaviors dependent on cerebellum-like structures. The odor categorization and VOR tasks both have input-output mappings that exhibit sharp nonlinearities and can be efficiently learned using sparse representations. In contrast, the forward modeling and feedback cancellation tasks have smooth input-output mappings and exhibit denser optima. These observations are consistent with our previous finding that more structured tasks favor denser coding levels than do random categorization tasks (Fig. 2E, F).

## Discussion

We have shown that the optimal granule cell coding level supporting cerebellar learning depends on the task being learned. While sparse representations are suitable for learning to categorize inputs into random categories, as predicted by classic theories, tasks involving structured input-output mappings benefit from much denser representations (Fig. 2). This reconciles such theories with the observation of dense granule cell activation during movement (Knogler et al., 2017; Wagner et al., 2017; Giovannucci et al., 2017; Badura and De Zeeuw, 2017; Wagner et al., 2019). We also show that, in contrast to the task-dependence of coding level, anatomical values of properties of granule cell and Purkinje cell connectivity are largely task-independent (Fig. 6). This distinction suggests that a stereotyped cerebellar architecture may support diverse representations optimized for diverse learning tasks.

### Relationship to previous theories

Previous studies assessed the learning performance of cerebellum-like systems with a model Purkinje cell that associates random patterns of mossy fiber activity with one of two randomly assigned categories (Marr, 1969; Albus, 1971; Brunel et al., 2004; Babadi and Sompolinsky, 2014; Litwin-Kumar et al., 2017; Cayco-Gajic et al., 2017), a common benchmark for artificial learning systems (Gerace et al., 2022). In this case, a low coding level increases the dimension of the granule cell representation, permitting more associations to be stored. The optimal coding level is low but not arbitrarily low, as extremely sparse representations introduce noise that hinders generalization (Barak et al., 2013; Babadi and Sompolinsky, 2014). A previous proposal also suggested that dense coding may improve generalization (Spanne and Jörntell, 2015), while our theory reveals that this property is task-dependent.

To examine a broader family of tasks, our learning problems extend previous studies in several ways. First, we consider inputs that may be constrained to a low-dimensional task subspace. Second, we consider input-output mappings beyond random categorization tasks. Finally, we assess generalization error for arbitrary locations on the task subspace, rather than only for noisy instances of previously presented inputs. As we have shown, these considerations require a complete analysis of the inductive bias of cerebellum-like networks (Fig. 4). Our analysis generalizes previous approaches (Barak et al., 2013; Babadi and Sompolinsky, 2014; Litwin-Kumar et al., 2017) that focused on dimension and noise alone. In particular, both dimension and noise for random patterns can be directly calculated from the kernel function (Fig. 3C; see Appendix).

Our theory builds upon techniques that been developed for understanding properties of kernel regression (Sollich, 1998; Jacot et al., 2018; Bordelon et al., 2020; Canatar et al., 2021b; Simon et al., 2021). Kernel approximations of wide neural networks are a major area of current research providing analytically tractable theories (Rahimi and Recht, 2007; Jacot et al., 2018; Chizat et al., 2018). Prior studies have analyzed kernels corresponding to networks with zero (Cho and Saul, 2010) or mean-zero Gaussian thresholds (Basri et al., 2019; Jacot et al., 2018), which in both cases produce networks with a coding level of 0.5. Ours is the first study of the effects of nonzero average thresholds. Our full characterization of the eigenvalue spectra and their decay rates as a function of the threshold extends previous work (Bach, 2017; Bietti and Bach, 2021). Furthermore, artificial neural network studies typically assume either fully-connected or convolutional layers, yet pruning connections after training barely degrades performance (Han et al., 2015; Zhang et al., 2018). Our results support the idea that sparsely connected networks may behave like dense ones if the representation is distributed (Fig. 5), providing insight into the regimes in which pruning preserves performance.

While the optimal coding level is task-dependent, optimal properties of the expansion connectivity are not (Fig. 6). Sparse connectivity and a large expansion ratio ensure uniform sampling of the task subspace (Fig. 5), reducing redundancy in the expansion layer representation. The resulting network has a kernel that approaches that of an infinitely wide network. These anatomical parameters are therefore necessary for producing a suitable representation without changing the inductive bias that is set by the coding level.

### Assumptions and extensions

We have made several simplifying assumptions in our model. First, when comparing the inductive biases of networks with different coding levels, we assumed that inputs were normalized and distributed uniformly in a linear subspace of the input layer activity. This allowed us to decompose the target function into a basis in which we could directly compare eigenvalues, and hence learning performance, for different coding levels (Fig. 4E-G). A similar analysis can be performed when inputs are not uniformly distributed, but in this case the basis is determined by an interplay between this distribution and the nonlinearity of expansion layer neurons, making the analysis more complex (see Appendix). A second assumption is that generalization is assessed for inputs drawn from the same distribution as used for learning. Recent and ongoing work on out-of-distribution generalization may permit relaxations of this assumption (Shen et al., 2021; Canatar et al., 2021a). Third, although we have modeled several timing-dependent tasks (Fig. 7), our learning rule does not exploit temporal information, and we assume that temporal dynamics of granule cell responses are largely inherited from mossy fibers. Finally, our theory assumes an infinitely wide expansion layer. When *P* is small enough that performance is limited by number of samples, this assumption is appropriate, but finite-size corrections to our theory are an interesting direction for future work.

We have also quantified coding level by the fraction of neurons, modeled as rate units, that are above firing threshold. We focused on coding levels *f* < 0.5, as extremely dense codes are rarely found in experiments (Olshausen and Field, 2004), but our theory applies for *f >* 0.5 as well. In general, representations with coding levels of *f* and 1 − *f* perform similarly in our model due to a symmetry in their associated eigenvalues (Supp. Fig. 2 and Appendix). Under the assumption that the energetic costs associated with neural activity are minimized, the *f* < 0.5 region is likely the biologically plausible one. Our qualitative results also do not depend on which threshold-nonlinear function (Supp. Fig. 4) or input dimension (Supp. Fig. 5, Supp. Fig. 1) we use.

Moreover, our results also hold when we assume granule cell spiking exhibits Poisson variability and quantify coding level as the fraction of neurons that have nonzero spike counts (Supp. Fig. 6; Fig. 7C). In general, increased spike count leads to improved performance as noise associated with spiking variability is reduced. Granule cells have been shown to exhibit reliable burst responses to mossy fiber stimulation (Chadderton et al., 2004), motivating models using deterministic responses or sub-Poisson spiking variability, but further work is needed to quantitatively compare variability in model and experiment.

### Implications for cerebellar representations

Our results predict that qualitative differences in the coding levels of cerebellum-like systems, across brain regions and/or across species, reflect an optimization to distinct tasks (Fig. 7). In the *Drosophila* mushroom body, which is required for associative learning of odor categories, random and sparse subsets of Kenyon cells are activated in response to odor stimulation, consistent with our model (Fig. 7B; Turner et al., 2008; Honegger et al., 2011; Lin et al., 2014). In a model of the electrosensory system of the electric fish, the inferred coding level of a model constrained by the properties of granule cells is similar to that which optimizes task performance (Fig. 7C). Within the cerebellar cortex, heterogeneity in granule cell firing has been observed across cerebellar lobules, associated with both differences in intrinsic properties (Heath et al., 2014) and mossy fiber input (Witter and De Zeeuw, 2015). It would be interesting to correlate such physiological heterogeneity with heterogeneity in function across the cerebellum. Our model predicts that regions involved in behaviors with substantial low-dimensional structure, for example smooth motor control tasks, may exhibit higher coding levels than regions involved in categorization or discrimination of high-dimensional stimuli.

Our model also raises the possibility that individual brain regions may exhibit different coding levels at different moments in time, depending on immediate behavioral or task demands. Multiple mechanisms could support the dynamic adjustment of coding level, including changes in mossy fiber input (Ozden et al., 2012), Golgi cell inhibition (Eccles et al., 1966; Palay and Chan-Palay, 1974), or unsupervised plasticity of mossy fiber-to-granule cell synapses (Schweighofer et al., 2001). The predictions of our model are not dependent on which of these mechanisms are operative. A recent study demonstrated that local synaptic inhibition by Golgi cells controls the spiking threshold and hence the population coding level of cerebellar granule cells in mice (Fleming et al., 2022). Further, the authors observed that granule cell responses to sensory stimuli are sparse when movement-related selectivity is controlled for. This suggests that dense movement-related activity and sparse sensory-evoked activity are not incompatible.

While our analysis makes clear qualitative predictions concerning comparisons between the optimal coding levels for different tasks, in some cases it is also possible to make quantitative predictions about the location of the optimum for a single task. Doing so requires determining the appropriate time interval over which to measure coding level, which depends on the integration time constant of the readout neuron. It also requires estimates of the firing rates and biophysical properties of the expansion layer neurons. In the electrosensory system, for which a well-calibrated model exists and the learning objective is well-characterized (Kennedy et al., 2014), we found that the coding level estimated based on the data is similar to that which optimizes performance (Fig. 7C).

If coding level is task-optimized, our model predicts that manipulating coding level artificially will diminish performance. In the *Drosophila* mushroom body, disrupting feedback inhibition from the GABAergic anterior paired lateral neuron onto Kenyon cells increases coding level and impairs odor discrimination (Lin et al., 2014). A recent study demonstrated that blocking inhibition from Golgi cells onto granule cells results in denser granule cell population activity and impairs performance on an eye-blink conditioning task (Fleming et al., 2022). These examples demonstrate that increasing coding level during sensory discrimination tasks, for which sparse activity is optimal, impairs performance. Our theory predicts that decreasing coding level during a task for which dense activity is optimal, such as smooth motor control, would also impair performance.

While dense activity has been taken as evidence against theories of combinatorial coding in cerebellar granule cells (Knogler et al., 2017; Wagner et al., 2019), our theory suggests that the two are not incompatible. Instead, the coding level of cerebellum-like regions may be determined by behavioral demands and the nature of the input to granule-like layers (Muscinelli et al., 2022). Sparse coding has also been cited as a key property of sensory representations in the cerebral cortex (Olshausen and Field, 1996). However, recent population recordings show that such regions exhibit dense movement-related activity (Musall et al., 2019), much like in cerebellum. While the theory presented in this study does not account for the highly structured recurrent interactions that characterize cerebrocortical regions, it is possible that these areas also operate using inductive biases that are shaped by coding level in a similar manner to our model.

## Acknowledgments

The authors thank L. F. Abbott, N. A. Cayco-Gajic, N. Sawtell, and S. Fusi for helpful discussions and comments on the manuscript. M. X. was supported by NIH grant T32-NS064929. S. M. was supported by the Simons and Swartz Foundations. K. D. H. was supported by a grant from the Washington Research Foundation. A. L.-K. was supported by the Simons and Burroughs Wellcome Foundations. M. X., S. M., and A. L.-K. were also supported by the Gatsby Charitable Foundation and NSF NeuroNex award DBI-1707398.

## Author contributions

M. X., K. D. H., and A. L.-K. conceived the study. All authors performed analytical calculations and simulations. M. X. prepared the first draft of the manuscript, and all authors edited the manuscript.

## Methods

### Network model

The expansion layer activity is given by

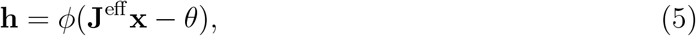

where **J**^eff^ = **JA** describes the selectivity of expansion layer neurons to task variables. For most simulations, **A** is an *N* × *D* matrix sampled with random, orthonormal columns and **J** is an *M* × *N* matrix with i.i.d. unit Gaussian entries. The nonlinearity *φ* is a rectified linear (ReLU) activation function *φ*(*x*) = max(*x*, 0) applied element-wise. The input layer activity is given by

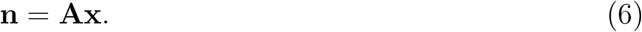

### Sparsely connected networks

To model sparse excitatory connectivity, we generated a sparse matrix **J**^*E*^, where each row contains precisely *K* nonzero elements at random locations. The nonzero elements are either identical and equal to 1 (homogeneous excitatory weights) or sampled from a unit truncated normal distribution (heterogeneous excitatory weights). To model global feedforward inhibition that balances excitation, **J** = **J**^*E*^ − **J**^*I*^, where **J**^*I*^ is a dense matrix with every element equal to 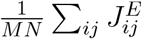.

For Fig. 5B, Fig. 6A, B, Fig. 7B, **J**, sparsely connected networks were generated with homogeneous excitatory weights and global inhibition. For Fig. 5E, the network with clustered representations was generated with homogeneous excitatory weights without global inhibition. For Fig. 5C, F, networks were generated with heterogeneous excitatory weights and global inhibition.

### Clustered representations

For clustered input-layer representations, each input layer neuron encodes one task variable (that is, **A** is a block matrix, with nonoverlapping blocks of *N/D* elements equal to 1 for each task variable). In this case, in order to obtain good performance, we found it necessary to fix the coding level for each input pattern, corresponding to winner-take-all inhibition across the expansion layer.

### Dimension

The dimension of the expansion layer representation (Fig. 3D) is given by (Abbott et al., 2011; Litwin-Kumar et al., 2017):

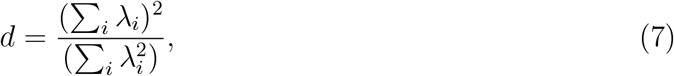

where *λ*_*i*_ are the eigenvalues of the covariance matrix 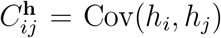 of expansion layer responses (not to be confused with *λ*_*α*_, the eigenvalues of the kernel operator). The covariance is computed by averaging over inputs **x**.

### Learning tasks

#### Random categorization task

In a random categorization task (Fig. 2E, Fig. 7B, Supp. Fig. 1, Supp. Fig. 3), the network learns to associate a random input pattern **x**^*µ*^ ∈ ℝ^*D*^ for *µ* = 1, …, *P* with a random binary category *y*^*µ*^ = ±1. The elements of **x**^*µ*^ are drawn i.i.d. from a normal distribution with mean 0 and variance 1*/D*. Test patterns 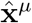^*µ*^ are generated by adding noise to the training patterns:

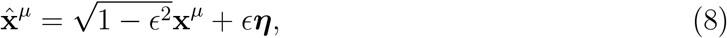

Where 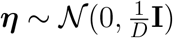. For Fig. 2E, Fig. 7B, and Supp. Fig. 3, we set *E* = 0.1.

#### Gaussian process tasks

To generate a family of tasks with continuously varying outputs (Fig. 2D,F, Fig. 4F,G, Fig. 5B, and Fig. 6), we sampled target functions from a Gaussian Process (GP; Rasmussen and Williams, 2006), *f* (**x**) ∼ 𝒢𝒫(0, *C*), with covariance

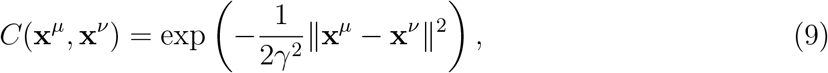

where *γ* determines the spatial scale of variations in *f*. Training and test patterns are drawn uniformly on the unit sphere.

#### Learning of readout weights

With the exception of the ELL task, we performed unregularized least squares regression to determine the readout weights **w**. For the ELL sensory cancellation task (Fig. 7C), we used *ℓ*^2^ regularization, a.k.a. ridge regression:

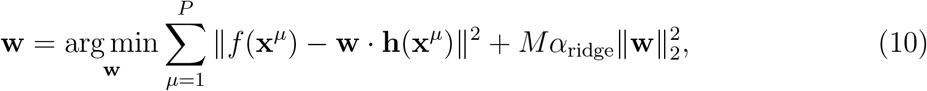

where *α*_ridge_ is the regularization parameter. Solutions were found using Python’s scikit-learn package (Pedregosa et al., 2011).

#### Performance metrics

For tasks with continuous targets, the prediction of the network is given by 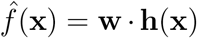, where **w** are the synaptic weights of the readout from the expansion layer. Error is measured as relative mean square error

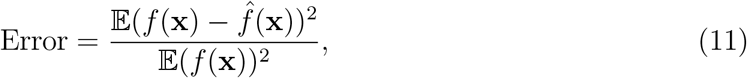

where in practice we use a large test set to estimate this error over **x** drawn from the distribution of test patterns. For categorization tasks, the network’s prediction is given by 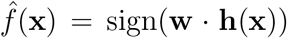. Performance is measured as the fraction of incorrect predictions. Error bars represent standard error of the mean across realizations of network weights and tasks.

#### Optimal granule–Purkinje cell weight distribution

We adapted our model to allow for comparisons with Brunel et al. (2004) by constraining readout weights **w** to be nonnegative and adding a bias, *f* (**x**) = **w**·**h**(**x**)+*b*. To guarantee that the target function is nonnegative, we set *f* (**x**) ∈ {0, 1} for the random categorization task and *f* (**x**) ← |*f* (**x**)| for the Gaussian process tasks. The weights and bias were determined with the Python convex optimization package cvxopt (Andersen et al., 2011).

### Model of two-joint arm

We implemented a biophysical model of a planar two-joint arm (Fagg et al., 1997). The state of the arm is specified by six variables: joint angles 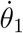 and 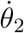, angular velocities *θ*?1 and *θ*?2, and torques *u*_1_ and *u*_2_. The upper and lower segments of the arm have lengths *l*_1_ and *l*_2_ and masses *m*_1_ and *m*_2_, respectively. The arm has the following dynamics:

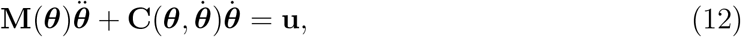

where **M**(***θ***) is the inertia matrix and 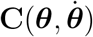 is the matrix of centrifugal, Coriolis, and friction forces:

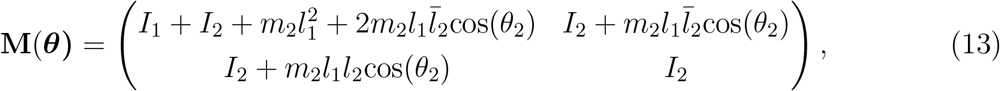

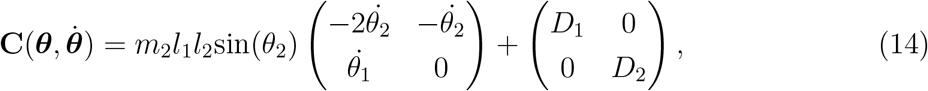

where 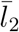 is the center of mass of the lower arm, *I*_1_ and *I*_2_ are moments of inertia and *D*_1_ and *D*_2_ are friction terms of the upper and lower arm respectively. These parameters were *m*_1_ = 3 kg, *m*_2_ = 2.5 kg, *l*_1_ = 0.3 m, *l*_2_ = 0.35 m, 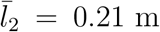, *I*_1_ = 0.1 kg m^2^, *I*_2_ = 0.12 kg m^2^, *D*_1_ = 0.05 kg m^2^ and *D*_2_ = 0.01 kg m^2^.

The task is to predict the position of the hand based on the forward dynamics of the two-joint arm system, given the arm initial condition and the applied torques. More precisely, the *P* network inputs **x**^*µ*^ were generated by sampling 6-dimensional Gaussian vectors with covariance matrix **C** = diag 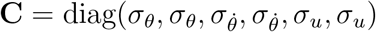, to account for the fact that angles, angular velocities and torques might vary on different scales across. For our results, we used 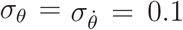 and *s*_*u*_ = 1. Each sample **x**^*µ*^ was then normalized and used to generate initial conditions of the arm, by setting 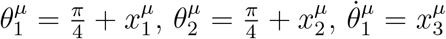, and 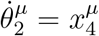. Torques were generated by setting 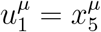 and 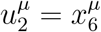. The target was constructed by running the dynamics of the arm forward in time for a time *δ* = 0.2 s, and by computing the difference in position of the “hand” (i.e. the end of the lower segment) in Cartesian coordinates. As a result, the target in this task is two-dimensional, with each target dimension corresponding the one of the two Cartesian coordinates of the hand. The overall performance is assessed by computing the error on each task separately and then averaging the errors.

### Model of *Drosophila* olfactory system

The model parameters are consistent with anatomical values for the *Drosophila* olfactory system ((Modi et al., 2020); Table 1). The input dimension *D* and *N* is based on the estimate of the number of antennal lobe glomeruli. The network learned a random categorization task.

**Table 1:** Summary of simulation parameters.

| Figure panel | Network parameters | Task parameters |
| --- | --- | --- |
| 2E | $M = 10,000$ | $D = 50, P = 1000, \epsilon = 0.1$ |
| 2F, 4G, 5B (full) | $M = 200,000$ | $D = 3, P = 30$ |
| 5B,E | $M = 200,000, N = 7000, K = 4$ | $D = 3, P = 30$ |
| 6A | $S = MK = 10,000, N = 100, f = 0.3$ | $D = 3, P = 200$ |
| 6B | $N = 700, K = 4, f = 0.3$ | $D = 3, P = 200$ |
| 6C | $M = 5000, f = 0.3$ | $D = 3, P = 100, \gamma = 1$ |
| 6D | $M = 1000$ | $D = 3, P = 50$ |
| 7A | $M = 20,000, dt = 0.01$ | $D = 6, P = 100, \delta = 20,$ |
| 7B | $M = 10,000, N = 50, K = 7$ | $D = 50, P = 100, \epsilon = 0.1$ |
| 7C | $M = 20,000$ | $N_{\text{trials}} = 4, \alpha_{\text{ridge}} = 0.01$ |
| 7D | $M = 20,000$ | $D = 1, P = 30$ |
| Suppl. 1A,B | $M = 10,000$ | $D = 50$ |
| Suppl. 4 | $M = 20,000$ | $D = 3, P = 30$ |
| Suppl. 5 | $M = 20,000$ | $P = 30, \gamma = 2$ |
| Suppl. 6 | $M = 20,000$ | $D = 3, P = 200, \gamma = 2$ |
| Suppl. 7 | $M = 20,000$ | $D = 3$ |

### Model of electrosensory lateral line lobe (ELL)

We simulated 20,000 granule cells using the biophysical model of Kennedy et al. (2014). We varied the granule cell layer coding level by adjusting the spiking threshold parameter in the model. For each choice of threshold, we generated 30 different trials of spike rasters. Each trial is 160 ms long with a 1 ms time bin and consists of a time-locked response to an electric organ discharge command. Trial-to-trial variability in the model granule cell responses arises from noise in the mossy fiber responses. To generate training and testing data, we sampled 4 trials (*P* = 640 patterns) from the 30 total trials for training and 10 trials for testing (1600 patterns). Coding level is measured as the fraction of granule cells that spike at least once in the training data. We repeated this sampling process 30 times.

The targets were smoothed broad-spike responses of 15 MG cells time-locked to an electric organ discharge command measured during experiments (Muller et al., 2019). The original data set consisted of 55 MG cells, each with a 300 ms long spike raster with a 1 ms time bin. The spike rasters were trial-averaged and then smoothed with a Gaussian-weighted moving average with a 10 ms time window. Only MG cells whose maximum spiking probability across all time bins exceeded 0.01 after smoothing were included in the task. The same MG cell responses were used for both training and testing. To match the length of the granule cell data, we discarded MG cell data beyond 160 ms and then concatenated 4 copies of the 160 ms long responses for training and 10 copies for testing. We measured the ability of the model to construct MG cell targets out of granule cell activity, generalizing across noise in granule cell responses. Errors for each MG cell target were averaged across the 30 repetitions of sampling of training and testing data, and then averaged across targets. Standard error of the mean was computed across the 30 repetitions.

### Model of vestibulo-ocular reflex (VOR)

Recordings of Purkinje cell activity in monkeys suggest that these neurons exhibit piecewise-linear tuning to head velocity (Lisberger et al., 1994). Thus, we designed piecewise-linear target functions with head velocity *v* as a one-dimensional input:

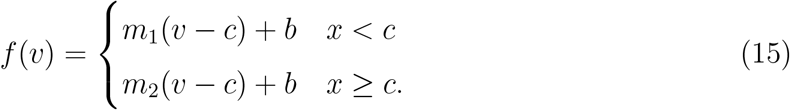

Inputs *v* were sampled uniformly from [−1, 1] 100 times. We generated 25 total target functions using all combinations of slopes *m*_1_ and *m*_2_ sampled from 5 equally spaced points on the interval [−2, 2]. We set *b* = 0.1 and *c* = −0.2.

Mossy fiber responses to head velocity input were modeled as exponential tuning curves:

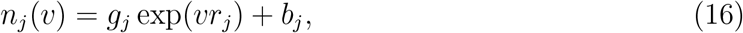

where *g*_*j*_ is a gain term, *r*_*j*_ ∈ ±1 determines a mossy fiber preference for positive or negative velocities, and *b*_*j*_ is the baseline firing rate. We generated 24 different tuning curves from all combinations of the following parameter values: The gain *g*_*j*_ was sampled from 6 equally spaced points on the interval [0.1, 1], *r*_*j*_ was set to either −1 or 1, and *b*_*j*_ was set to either 0 or 1. Qualitative results did not depend strongly on this parameterization. Errors were averaged across targets.

