## Supplemental Figures for "Task-dependent optimal representations for cerebellar learning"

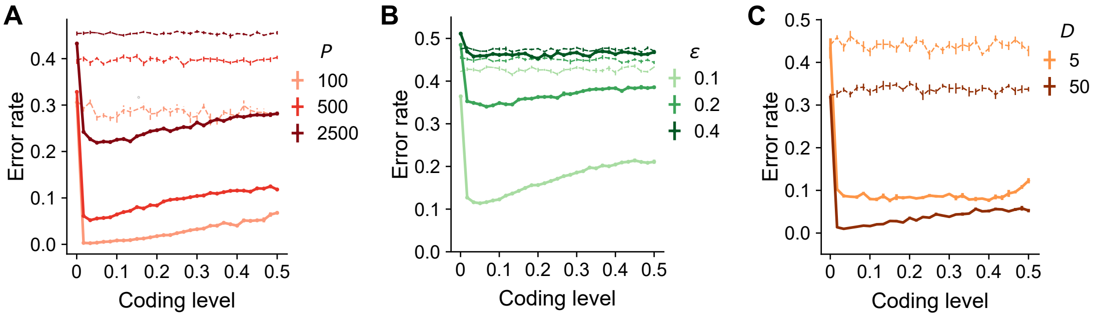


**Supplemental Figure 1: Sparse coding levels are sufficient for random categorization tasks irrespective of number of samples, noise level, and dimension.**
A) Error as a function of coding level for networks trained to perform random categorization tasks (as in Fig. 2E but with a wider range of associations $P$). Performance is measured for noisy instances of previously seen inputs. $D = 50$. Dashed lines indicate the performance of a readout of the input layer. Standard error of the mean was computed across 20 realizations of network weights and tasks.
B) Same as in (A) but fixing the number of associations and varying the noise $\epsilon$ which controls the deviation of test patterns from training patterns. $D = 50$.
C) Same as in (A) but varying the input dimension $D$. To improve performance for small $D$, we fixed the coding level for each pattern. $P = 200, \epsilon = 0.1$ For small $D$, the curve of error rate against coding level is more flat, but low coding levels are still sufficient to saturate performance.

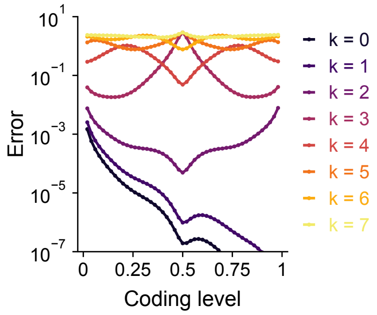


**Supplemental Figure 2: Error as a function of coding level for learning pure-frequency spherical harmonic functions.**
Frequency is indexed by $k$. Errors are calculated using analytically using Equation (4) and represent the predictions of the theory for an infinitely large expansion. Curves are symmetric around $f = 0.5$ except for $k = 0$ and $k = 1$. Results are shown for $D = 3$.

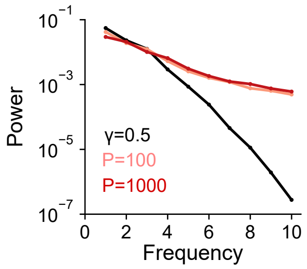


**Supplemental Figure 3: Frequency content of categorization tasks.**
Power as a function of frequency for random categorization tasks (colors) and for Gaussian process task (black). Power is averaged over realizations of target functions.

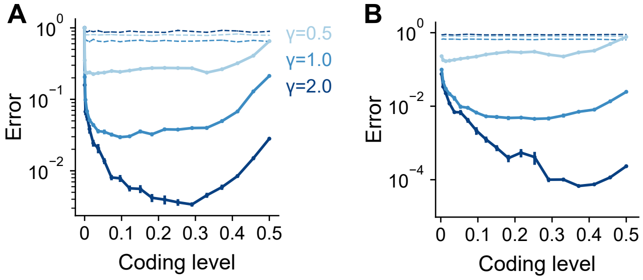


**Supplemental Figure 4: Task-dependence of optimal coding level is robust across activation functions.**
Error as a function of coding level for networks with A) Heaviside and B) rectified power-law (with power 2, $\phi(x) = [\max(0, x)]^2$) nonlinearity in the expansion layer. Networks learned Gaussian process targets. Dashed lines indicate the performance of a readout of the input layer. Standard error of the mean was computed across 10 realizations of network weights and tasks in (A) and 50 in (B). Parameters: $M = 20,000$, $P = 30$, $D = 3$.

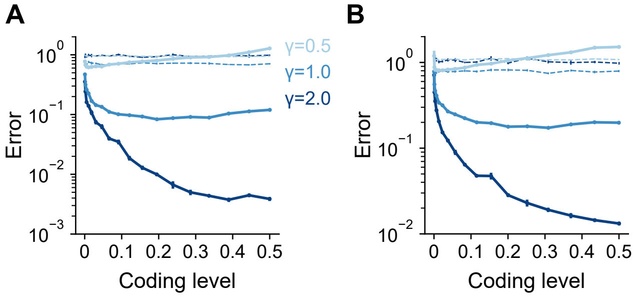


**Supplemental Figure 5: Task-dependence of optimal coding level is robust across input dimensions.**
Error as a function of coding level for networks learning Gaussian process targets with input dimension $D = 5$ (A) and $D = 7$ (B). Dashed lines indicate the performance of a readout of the input layer. Standard error of the mean was computed across 10 realizations of network weights and tasks. Parameters: $M = 20,000$, $P = 30$.

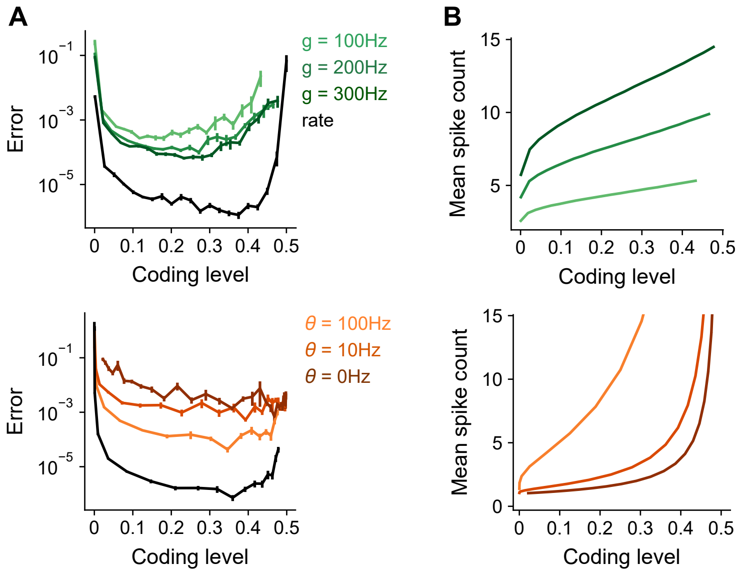


**Supplemental Figure 6: Optimal coding levels remain dense in the presence of spiking noise.**

A) Error as a function of coding level in a spiking model). The firing rate of neuron $i$ (in Hz) is given by $h_i^\mu = \phi(g\mathbf{J}_i^{\text{eff}}\mathbf{x}^\mu - \theta)$, where $g$ is a gain term that adjusts the amplitude of the activity and $\theta$ is the activation threshold. The spike count $s_i$ for a neuron $i$ in response to pattern $\mu$ is sampled from a Poisson distribution: $s_i = \text{Pois}(h_i^\mu \tau)$. $\tau$ represents the time window in which a Purkinje cell integrates spikes, and is set to 0.1 s. Coding level is measured as the fraction of cells with a nonzero spike count. Coding level is adjusted by tuning either the activation threshold $\theta$ (top) or the gain $g$ (bottom). Black curve shows the performance of a rate model as in the main text. Standard error of the mean was computed across 10 realizations of network weights.

B) Mean spike count of active expansion layer neurons during the time window $\tau$ as a function of coding level.

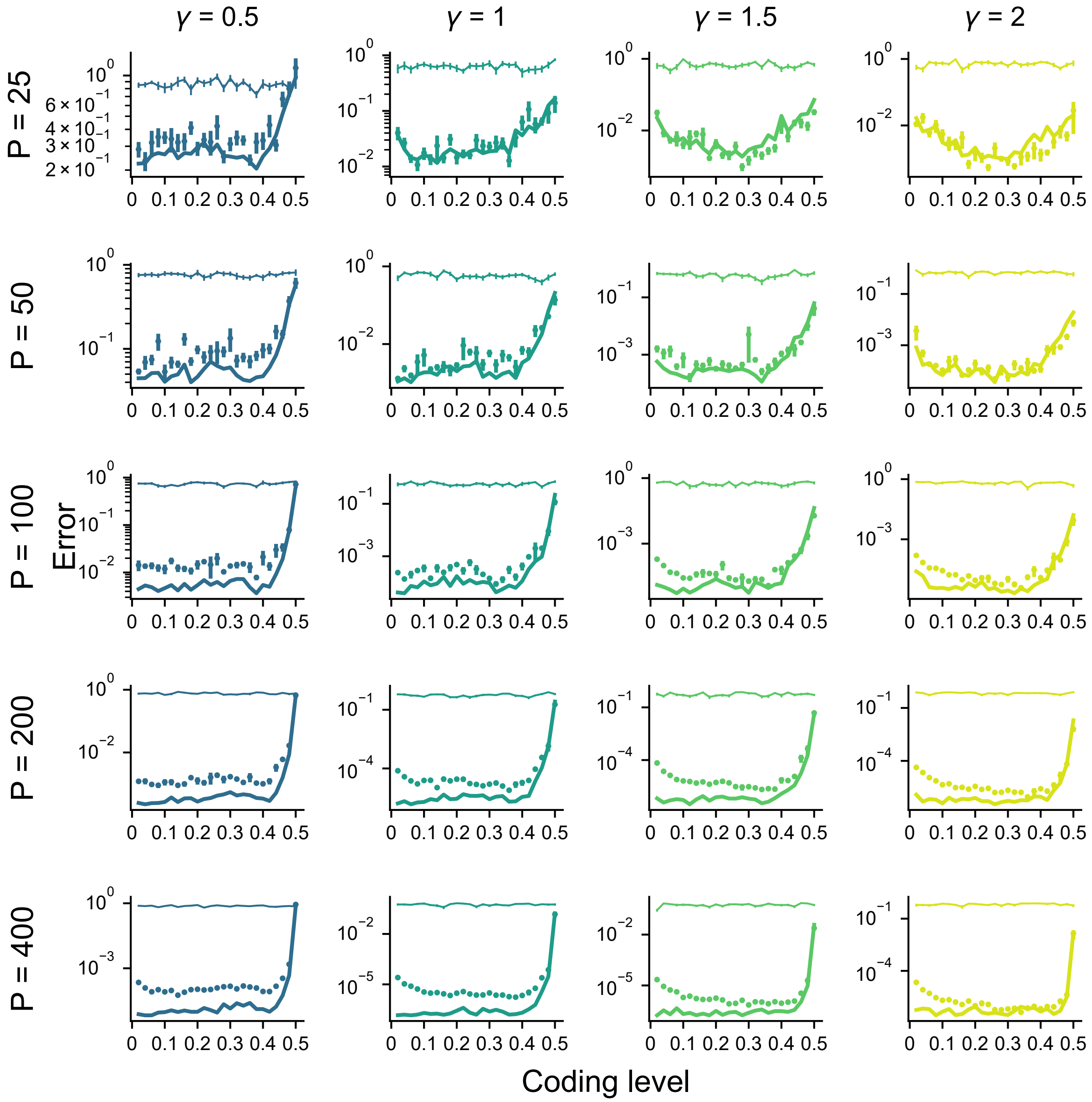


**Supplemental Figure 7: Error as a function of coding level across different values of $P$ and $\gamma$.**
Dots denote performance of a readout of the expansion layer in simulations. Thin lines denote performance of a readout of the input layer in simulations. Thick lines denote theory for expansion layer readout performance. Standard error of the mean was computed across 10 realizations of network weights and tasks. Parameters: $D = 3$, $M = 20{,}000$.
