## Appendix for "Task-dependent optimal representations for cerebellar learning"

### A Connection between kernel and previous theories

Previous theories (Babadi and Sompolinsky, 2014; Litwin-Kumar et al., 2017) studied generalization performance for random clusters of inputs associated with binary targets, where test patterns are formed by adding noise to training patterns (Fig. 4A). The readout is trained using a supervised Hebbian rule with mean-subtracted expansion layer responses, $\mathbf{w} = \sum_\mu y^\mu(\mathbf{h}^\mu - \bar{\mathbf{h}})$, with $\bar{\mathbf{h}} = \frac{1}{P}\sum_{\mu=1}^{P} \mathbf{h}^\mu$. The net input to a readout in response to a test pattern $\hat{\mathbf{h}}^\mu$ from cluster $\mu$ is $g^\mu = \mathbf{w} \cdot (\hat{\mathbf{h}}^\mu - \bar{\mathbf{h}})$. The statistics of $g^\mu$ determine generalization performance. For a Hebbian readout, the error rate is expressed in terms of the signal-to-noise ratio (SNR) (Babadi and Sompolinsky, 2014):

$$P(\text{Error}) = \frac{1}{2}\,\text{erfc}\left(\sqrt{\text{SNR}/2}\right). \tag{17}$$

SNR is given in terms of the mean and variance of $g^\mu$:

$$\text{SNR} = \frac{\left(\mathbb{E}_\mu\left[y^\mu g^\mu\right]\right)^2}{\text{Var}\left(g^\mu\right)}. \tag{18}$$

The numerator of SNR is proportional to the average overlap of the expansion layer representations of training and test patterns belonging to the same cluster, which can be expressed in terms of the kernel function $K$:

$$\mathop{\mathbb{E}}_{\mu}\left[y^\mu g^\mu\right] = \mathop{\mathbb{E}}_{\mu}\left[(\hat{\mathbf{h}}^\mu - \bar{\mathbf{h}}) \cdot (\mathbf{h}^\mu - \bar{\mathbf{h}})\right] = M \mathop{\mathbb{E}}_{\mu}\left[K(\hat{\mathbf{x}}^\mu, \mathbf{x}^\mu)\right] - \bar{\mathbf{h}} \cdot \bar{\mathbf{h}}. \tag{19}$$

For large networks with Gaussian i.i.d. expansion weights, $K(\hat{\mathbf{x}}^\mu, \mathbf{x}^\mu) = K(t)$, where $t = \hat{\mathbf{x}}^\mu \cdot \mathbf{x}^\mu$, and the above equation reduces to $MK(t_{\text{train/test}}) - \bar{\mathbf{h}} \cdot \bar{\mathbf{h}}$, where $t_{\text{train/test}}$ is the typical overlap of training and test patterns belonging to the same cluster. When $\mathbf{x}^\mu \cdot \mathbf{x}^\mu = 1$, $t_{\text{train/test}}$ can be written as $t_{\text{train/test}} = 1 - \Delta$, where $\Delta$ is a measure of within-cluster noise (Babadi and Sompolinsky, 2014; Litwin-Kumar et al., 2017).

Babadi and Sompolinsky (2014) demonstrated that, for random categorization tasks and when $M$ and $D$ are large, $\text{Var}\left(g^\mu\right) = C(\frac{1}{M} + Q^2\frac{1}{D})$ where $C$ is a constant and $Q \in [0, 1]$ is given by

$$Q^2 = \frac{\frac{1}{Z_h}\,\mathbb{E}_{\mu \neq \nu}\left[((\mathbf{h}^\mu - \bar{\mathbf{h}}) \cdot (\mathbf{h}^\nu - \bar{\mathbf{h}}))^2\right]}{\frac{1}{Z_x}\,\mathbb{E}_{\mu \neq \nu}\left[(\mathbf{x}^\mu \cdot \mathbf{x}^\nu)^2\right]}, \tag{20}$$

assuming the entries of $\mathbf{x}$ are zero-mean. $Z_a = \mathbb{E}_\mu[\|\mathbf{a}^\mu - \bar{\mathbf{a}}\|^2]$ for $a \in \{h, x\}$ normalizes the overlaps to the typical overlap of a pattern with itself. The quantity $Q^2$ is the ratio of the variance of overlaps between patterns belonging to different clusters in the expansion layer to that of the input layer. This describes the extent to which the geometry of the input layer representation is preserved in the expansion layer. When overlaps in the input layer are small, as they are for random clusters, $\frac{1}{Z_h}(\mathbf{h}^\mu - \bar{\mathbf{h}}) \cdot (\mathbf{h}^\nu - \bar{\mathbf{h}}) \approx \frac{Q}{Z_x} \cdot (\mathbf{x}^\mu \cdot \mathbf{x}^\nu)$ as $M \to \infty$. This relation illustrates that, for random clusters and $M \to \infty$, $Q$ is equal to the slope of the normalized kernel function $K(t)$ evaluated at $t = 0$. Litwin-Kumar et al. (2017) also showed that the dimension of the expansion layer representation is equal to $\frac{C'}{(\frac{1}{M} + Q^2 \frac{1}{D})}$, where $C'$ is a constant.

Thus, for the random categorization task studied in Babadi and Sompolinsky (2014); Litwin-Kumar et al. (2017), dimension and readout SNR can be calculated by evaluating $K(t_{\text{train/test}})$ and the slope of $K(t)$ at $t = 0$.

### B Dot-product kernels with arbitrary threshold

As $M \to \infty$, the normalized dot product between features (2) converges pointwise to

$$K(\mathbf{x}, \mathbf{x}') = \mathop{\mathbb{E}}_{\mathbf{J}_1} \left[ \phi(\mathbf{J}_1^T \mathbf{x} - \theta) \phi(\mathbf{J}_1^T \mathbf{x}' - \theta) \right], \tag{21}$$

where $\mathbf{J}_1$ is a row of the weight matrix $\mathbf{J}$ (without loss of generality, the first row) with entries drawn i.i.d. from a Gaussian distribution $\mathcal{N}(0, 1)$. Our goal is to compute (21) for a given $\theta$ and inputs drawn on the unit sphere $\mathbf{x}, \mathbf{x}' \in \mathbb{S}^{D-1}$.

Because the Gaussian weight distribution is spherically symmetric, (21) restricted to the unit sphere *for any nonlinearity* is only a function of the dot-product $t := \mathbf{x}^T \mathbf{x}'$, making the kernel a dot-product kernel $K(\mathbf{x}, \mathbf{x}') = K(t)$.

Denote by $J_i$ the entries of $\mathbf{J}_1$. Let $I_1 = \sum_{i=1}^{D} J_i x_i$ and $I_2 = \sum_{i=1}^{D} J_i x'_i$ be the pre-activations for each input. Then $(I_1, I_2)$ are jointly Gaussian with mean 0, variance 1, and covariance $\mathbb{E}[I_1 I_2] = t$. If $t > 0$, we can re-parameterize these pre-activations as the sum of an independent and shared component $I_i = y_i \sqrt{1-t} + z\sqrt{t}$, where $y_i \sim \mathcal{N}(0, 1)$ for $i = 1, 2$ and

$z \sim \mathcal{N}(0,1)$. In these coordinates, (21) becomes

$$\begin{aligned} K(t) &= \mathop{\mathbb{E}}_{y_1,y_2,z}\left[\phi(y_1\sqrt{1-t}+z\sqrt{t}-\theta)\phi(y_2\sqrt{1-t}+z\sqrt{t}-\theta)\right] \\ &= \mathop{\mathbb{E}}_{z}\left[\mathop{\mathbb{E}}_{y_1}[\phi(y_1\sqrt{1-t}+z\sqrt{t}-\theta)|z]\ \mathop{\mathbb{E}}_{y_2}[\phi(y_2\sqrt{1-t}+z\sqrt{t}-\theta)|z]\right] \\ &= \mathop{\mathbb{E}}_{z}\left[\mathop{\mathbb{E}}_{y_1}[\phi(y_1\sqrt{1-t}+z\sqrt{t}-\theta)|z]^2\right], \end{aligned} \tag{22}$$

where the second line follows from the conditional independence of $h_1|z$ and $h_2|z$ and the third from the fact that they are identically distributed. Similarly, if $t<0$, we can write $I_1 = y_1\sqrt{1-t}+z\sqrt{t}$, $I_2 = y_2\sqrt{1-t}-z\sqrt{t}$.

We will use (22) to solve for the kernel assuming $\phi$ is a ReLU nonlinearity. Let

$$g_1(t,z) = \mathbb{E}[\phi(y_1\sqrt{1-t}+z\sqrt{t}-\theta)|z]. \tag{23}$$

Using the fact that $\phi$ is nonzero only when $y_1\sqrt{1-t}+z\sqrt{t}-\theta>0$, i.e. for $y_1 > T = \frac{\theta - z\sqrt{t}}{\sqrt{1-t}}$, we obtain

$$\begin{aligned} g_1(t,z) &= (2\pi)^{-1/2}\int_T^\infty (y_1\sqrt{1-t}+z\sqrt{t}-\theta)e^{-y_1^2/2}\mathrm{d}y_1 \\ &= \left(\frac{1-t}{2\pi}\right)^{1/2} e^{-T^2/2} + \left(\frac{z\sqrt{t}-\theta}{2}\right)\operatorname{erfc}(T/\sqrt{2}). \end{aligned} \tag{24}$$

Performing a similar calculation for $t<0$ and collecting the results leads to:

$$K(t) = \begin{cases} \mathbb{E}_z\left[g_1(t,z)^2\right] & t>0 \\ \mathbb{E}_z\left[g_1(|t|,z)g_2(|t|,z)\right] & t<0 \end{cases}, \tag{25}$$

$$g_1(t,z) = \left(\frac{1-t}{2\pi}\right)^{1/2} e^{-T^2/2} + \left(\frac{z\sqrt{t}-\theta}{2}\right)\operatorname{erfc}(T_1/\sqrt{2}) \tag{26}$$

$$g_2(t,z) = \left(\frac{1-t}{2\pi}\right)^{1/2} e^{-T^2/2} + \left(\frac{-z\sqrt{t}-\theta}{2}\right)\operatorname{erfc}(T_2/\sqrt{2}) \tag{27}$$

$$T_1 = \frac{\theta - z\sqrt{t}}{\sqrt{1-t}}, \quad \text{and} \quad T_2 = \frac{\theta + z\sqrt{t}}{\sqrt{1-t}}. \tag{28}$$

### C Spherical harmonic decompositions

Our theory of generalization requires us to work in function spaces which are natural to the problem. The spherical harmonics are the natural basis for working with dot-product kernels on the sphere. For a thorough treatment of spherical harmonics, see Atkinson and Han (2012), whose notation we generally follow. Both our kernel and Gaussian process (GP) tasks are defined over the sphere in $D$ dimensions

$$\mathbb{S}^{D-1} = \{\mathbf{x} \in \mathbb{R}^D : \|\mathbf{x}\|_2 = 1\}. \tag{29}$$

A spherical harmonic $Y_{km}(\cdot)$—where $k$ indexes frequency and $m$ indexes modes of the same frequency—is a harmonic homogeneous polynomial of degree $k$ restricted to the sphere $\mathbb{S}^{D-1}$. For each frequency $k \in \mathbb{Z}$, there are $N(D,k)$ linearly independent polynomials, where

$$N(D,k) = \frac{2k+D-2}{k}\binom{k+D-3}{k-1}. \tag{30}$$

#### C.1 Decomposition of the kernel and target function

We remind the reader here of the setting for our theory:

1. Ridge regression using random features with a dot-product limiting kernel.
2. Data drawn uniformly from the unit sphere.

Let $\sigma$ be the Lebesgue measure on $\mathbb{S}^{D-1}$. We will denote the surface area of the sphere as

$$|\mathbb{S}^{D-1}| = \int_{\mathbb{S}^{D-1}} \mathrm{d}\sigma = \frac{2\pi^{D/2}}{\Gamma(D/2)}. \tag{31}$$

On the other hand, the uniform probability measure on the sphere, denoted by $\bar{\sigma}$, must integrate to 1, so $\bar{\sigma} = \sigma / |\mathbb{S}^{D-1}|$. Finally, we define the space of real-valued square integrable functions $L^2(\sigma)$ as the Hilbert space with inner product

$$\langle f, g\rangle_{L^2(\sigma)} = \int_{\mathbb{S}^{D-1}} f(\mathbf{x}) g(\mathbf{x})\, \mathrm{d}\sigma(\mathbf{x}), \tag{32}$$

and $\|f\|_{L^2(\sigma)} = \langle f, f\rangle_{L^2(\sigma)}^{1/2}$. The space $L^2(\bar{\sigma})$ is defined analogously.

Eigendecompositions describe the action of linear operators, not functions, thus we must associate a linear operator with our kernel for its eigenvalues to make sense. The kernel eigenvalues $\lambda_\alpha$ that we will use to compute the error in (4) are the eigenvalues of the integral operator $\mathcal{T}_K : L^2(\bar{\sigma}) \to L^2(\bar{\sigma})$ defined as

$$(\mathcal{T}_K f)(\mathbf{x}) = \langle K(\mathbf{x}, \cdot), f(\cdot) \rangle_{L^2(\bar{\sigma})} = \int_{\mathbb{S}^{D-1}} K(\mathbf{x}, \mathbf{x}') f(\mathbf{x}') \mathrm{d}\bar{\sigma}(\mathbf{x}'). \tag{33}$$

This is because $\bar{\sigma}$ is the data distribution, and these eigenvalues are approximated by the eigenvalues of the kernel matrix evaluated on a large but finite dataset (Koltchinskii and Giné, 2000). Similarly, we define the analogous operator $\mathcal{U}_K : L^2(\sigma) \to L^2(\sigma)$ under the measure $\sigma$ with eigenvalues $\xi_\alpha$. Since $\mathcal{T}_K = \mathcal{U}_K / |\mathbb{S}^{D-1}|$, the eigenvalues are related by

$$\lambda_\alpha = \frac{\xi_\alpha}{|\mathbb{S}^{D-1}|}, \tag{34}$$

and they share the same eigenfunctions, up to normalization. For the rest of this section we will study eigendecompositions of operator $\mathcal{U}_K$, which may be translated into statements about $\mathcal{T}_K$ via (34).[1]

Under mild technical conditions that our kernels satisfy, Mercer's theorem states that positive semidefinite kernels can be expanded as a series in the orthonormal basis of eigenfunctions $\psi_\alpha$ weighted by nonnegative eigenvalues $\xi_\alpha$:

$$K(\mathbf{x}, \mathbf{x}') = \sum_\alpha \xi_\alpha \psi_\alpha(\mathbf{x}) \psi_\alpha(\mathbf{x}'). \tag{35}$$

Again, $(\lambda_\alpha, \psi_\alpha)$ are eigenpairs for the operator $\mathcal{U}_K$ and form an orthonormal set under the $L^2(\sigma)$ inner product.

As stated earlier, the kernel (21) is spherically symmetric and thus a dot-product kernel. Because of this, we can take the eigenfunctions $\psi_\alpha$ to be the spherical harmonics $Y_{km}$. The index $\alpha$ is a multi-index into mode $m$ of frequency $k$. Writing the Mercer decomposition in the spherical harmonic basis gives:

$$K(\mathbf{x}, \mathbf{x}) = \sum_{k=0}^{\infty} \xi_k \sum_{m=1}^{N(D,k)} Y_{km}(\mathbf{x}) Y_{km}(\mathbf{x}'). \tag{36}$$

Because our kernel is rotation invariant, all $N(D, k)$ harmonics of frequency $k$ share eigen-

[1] These differences are liable to cause some confusion and pain when reading the literature.

value $\xi_k$.

Any function in $L^2(\sigma)$ can be expanded in the spherical harmonic basis as follows:

$$f(\mathbf{x}) = \sum_{k=0}^{\infty} \sum_{m=1}^{N(D,k)} c_{km} Y_{km}(\mathbf{x}), \text{ with } c_{km} = \langle f, Y_{km} \rangle_{L^2(\sigma)}. \tag{37}$$

The expansion is analogous to that of the Fourier series. In fact when $D = 2$, the spherical harmonics are sines and cosines on the unit circle.

#### C.2 Ultraspherical polynomials

Adding together all harmonics of a given frequency relates them to a polynomial in $t$ by the addition formula

$$\sum_{m=1}^{N(D,k)} Y_{km}(\mathbf{x}) Y_{km}(\mathbf{x}') = \frac{N(D,k)}{|\mathbb{S}^{D-1}|} P_{k,D}(\mathbf{x}^T \mathbf{x}'). \tag{38}$$

The polynomial $P_{k,D}(t)$ is the $k$th ultraspherical polynomial. These are also called Legendre or Gegenbauer polynomials, although these usually have different normalizations and can be defined more generally.

The ultraspherical polynomials $\{P_{k,D}\}$ form an orthogonal basis for

$$L^2([-1,1], (1-t^2)^{(D-3)/2} \mathrm{d}t).$$

As special cases, $P_{k,2}(t)$ and $P_{k,3}(t)$ are the classical Chebyshev and Legendre polynomials, respectively. For any $D$, the first two of these polynomials are $P_0(t) = 1$ and $P_1(t) = t$. We use the Rodrigues formula, which holds for $k \geq 0$ and $D \geq 2$, to generate these polynomials

$$P_{k,D}(t) = (-1/2)^k \frac{\Gamma((D-1)/2)}{\Gamma(k + (D-1)/2)} (1-t^2)^{(3-D)/2} \left( \frac{\mathrm{d}}{\mathrm{d}t} \right)^k (1-t^2)^{k+(D-3)/2}. \tag{39}$$

Combining (36) with the addition formula (38), we can express the kernel in terms of ultraspherical polynomials evaluated at the dot-product of the inputs:

$$K(t) = \sum_{k=0}^{\infty} \xi_k \frac{N(D,k)}{|\mathbb{S}^{D-1}|} P_{k,D}(t). \tag{40}$$

#### C.3 Computing kernel eigenvalues

The Funk-Hecke theorem states that

$$\int_{\mathbf{x}\in\mathbb{S}^{D-1}} K(\mathbf{x}^T\mathbf{x}')Y_k(\mathbf{x}')\mathrm{d}\sigma(\mathbf{x}') = |\mathbb{S}^{D-2}|Y_k(\mathbf{x})\int_{-1}^{1} K(t)P_{k,D}(t)(1-t^2)^{(D-3)/2}\mathrm{d}t. \tag{41}$$

Equation (41) implies that the eigenvalues of $\mathcal{U}_K$ are given as

$$\xi_k = |\mathbb{S}^{D-2}|\int_{-1}^{1} K(t)P_{k,D}(t)(1-t^2)^{(D-3)/2}\mathrm{d}t. \tag{42}$$

For our kernels, the kernel eigenvalues can be conveniently computed using polar coordinates. When the entries of $\mathbf{J}_1$ are i.i.d. unit Gaussian,

$$\begin{aligned}K(t) &= \int_{\mathbb{R}^D} \phi(\mathbf{J}_1^T\mathbf{x}-\theta)\phi(\mathbf{J}_1^T\mathbf{x}'-\theta)(2\pi)^{-D/2}e^{-\|\mathbf{J}_1\|^2/2}\,\mathrm{d}(\mathbf{J}_1)_1\cdots\mathrm{d}(\mathbf{J}_1)_D \\ &= (2\pi)^{-D/2}\int_0^\infty e^{-r^2/2}r^{D-1}\int_{\mathbb{S}^{D-1}} \phi(r\hat{\mathbf{J}}^T\mathbf{x}-\theta)\phi(r\hat{\mathbf{J}}^T\mathbf{x}'-\theta)\mathrm{d}\sigma(\hat{\mathbf{J}})\mathrm{d}r,\end{aligned}$$

where $\hat{\mathbf{J}} = \mathbf{J}_1/r$ and $r = \|\mathbf{J}_1\|$. The ReLU nonlinearity is positively homogeneous, so $\phi(r\hat{\mathbf{J}}^T\mathbf{x}-\theta) = r\phi(\hat{\mathbf{J}}^T\mathbf{x}-\theta/r)$. We can write

$$\begin{aligned}K(t) &= (2\pi)^{-D/2}\int_0^\infty e^{-r^2/2}r^{D+1}\underbrace{\int_{\mathbb{S}^{D-1}}(\hat{\mathbf{J}}^T\mathbf{x}-\theta/r)_+(\hat{\mathbf{J}}^T\mathbf{x}'-\theta/r)_+\mathrm{d}\sigma(\hat{\mathbf{J}})}_{:=K_{\text{shell}}(t;\theta/r)}\,\mathrm{d}r \\ &= (2\pi)^{-D/2}\int_0^\infty e^{-r^2/2}r^{D+1}K_{\text{shell}}(t;\theta/r)\mathrm{d}r,\end{aligned} \tag{43}$$

where we have introduced a new kernel $K_{\text{shell}}(t;\theta)$ which is $|\mathbb{S}^{D-1}|$ times the dot-product kernel that arises when the weights are distributed uniformly on the sphere ($\sigma$ is not the probability measure). The above equation shows that the network restricted to inputs $\mathbf{x},\mathbf{x}'\in\mathbb{S}^{D-1}$ has different kernels depending on whether the weights are sampled according to a Gaussian distribution or uniformly on the sphere. Without the threshold, this difference disappears due to the positive homogeneity of the ReLU (Cho and Saul, 2010).

Next we expand the nonlinearity in the spherical harmonic basis (following Bietti and Bach,

2021; Bach, 2017)

$$\phi(\hat{\mathbf{J}}^T\mathbf{x}-\theta) = (\hat{\mathbf{J}}^T\mathbf{x}-\theta)_+ = \sum_{k=0}^{\infty} a_k(\theta) \sum_{j=1}^{N(D,k)} Y_{kj}(\hat{\mathbf{J}})Y_{kj}(\mathbf{x}), \tag{44}$$

where the $k$th coefficient is given by the Funk-Hecke formula (41) as

$$a_k(\theta) = |\mathbb{S}^{D-2}| \int_{-1}^{1} (t-\theta)_+ P_k(t)(1-t^2)^{(D-3)/2}\mathrm{d}t, \tag{45}$$

and we explicitly note the dependence on $\theta$. Using the representation (44), we can recover the eigendecomposition:

$$\begin{aligned}
K_{\text{shell}}(t;\theta) &= \int_{\mathbb{S}^{D-1}} (\hat{\mathbf{J}}^T\mathbf{x}-\theta)_+(\hat{\mathbf{J}}^T\mathbf{x}'-\theta)_+\mathrm{d}\sigma(\hat{\mathbf{J}}) \\
&= \sum_{k,k'} a_k(\theta)a_{k'}(\theta) \sum_{j,j'} Y_{kj}(\mathbf{x})Y_{k'j'}(\mathbf{x}') \underbrace{\int_{\mathbb{S}^{D-1}} Y_{kj}(\hat{\mathbf{J}})Y_{k'j'}(\hat{\mathbf{J}})\mathrm{d}\sigma(\hat{\mathbf{J}})}_{\delta_{kk'}\delta_{jj'}} \\
&= \sum_{k} a_k(\theta)^2 \sum_{j} Y_{kj}(\mathbf{x})Y_{kj}(\mathbf{x}') \\
&= \sum_{k} a_k(\theta)^2 \frac{N(k,D)}{|\mathbb{S}^{D-1}|} P_k(t),
\end{aligned} \tag{46}$$

which follows from orthonormality and the addition formula (38). We have that $a_k(\theta)^2$ is the $k$th eigenvalue of $K_{\text{shell}}(t;\theta)$.

Using (46) in (43) leads to

$$K(t) = \sum_{k} \frac{N(k,D)}{|\mathbb{S}^{D-1}|} P_k(t)(2\pi)^{-D/2} \int_0^{\infty} e^{-r^2/2} r^{D+1} a_k(\theta/r)^2 \mathrm{d}r, \tag{47}$$

i.e. the eigenvalues satisfy

$$\xi_k = (2\pi)^{-D/2} \int_0^{\infty} e^{-r^2/2} r^{D+1} a_k(\theta/r)^2 \mathrm{d}r. \tag{48}$$

##### C.3.1 Eigenvalues of $K_{\text{shell}}$

It is possible to compute $a_k(\theta)$ analytically (Bietti and Bach, 2021; Bach, 2017). Letting

$$I_{\alpha,k}(\theta) = \int_\theta^1 t^\alpha P_k(t)(1-t^2)^{(D-3)/2}\mathrm{d}t, \tag{49}$$

we have that (45) reduces to $a_k(\theta) = |\mathbb{S}^{D-2}|\,(I_{1,k}(\theta^*) - \theta I_{0,k}(\theta^*))$. Equation (49) requires $\theta \in [-1,1]$, but $\theta/r \to \pm\infty$ in (48) as $r \to 0$. So we take $\theta^* = \min(\max(\theta,-1),1)$, which still assures that (45) is satisfied. For the rest of this section, assume wlog that $\theta \in [-1,1)$.

Using Rodrigues' formula (39) in (49) gives

$$\begin{aligned}
I_{\alpha,k}(\theta) &= \underbrace{(-1/2)^k \frac{\Gamma((D-1)/2)}{\Gamma(k+(D-1)/2)}}_{:=C} \int_\theta^1 t^\alpha \left(\frac{\mathrm{d}}{\mathrm{d}t}\right)^k (1-t^2)^{k+(D-3)/2}\mathrm{d}t \\
&= C\int_\theta^1 t^\alpha \left(\frac{\mathrm{d}}{\mathrm{d}t}\right)^k (1-t^2)^{k+(D-3)/2}\mathrm{d}t
\end{aligned}$$

which may be integrated by parts. We will treat $\alpha = 0$ and 1 separately.

In the case of $\alpha = 0$, since $t^\alpha = 1$ we have the integral of a derivative, so for $k \geq 1$

$$\begin{aligned}
I_{0,k}(\theta) &= C\int_\theta^1 \left(\frac{\mathrm{d}}{\mathrm{d}t}\right)^k (1-t^2)^{k+(D-3)/2}\mathrm{d}t \\
&= C\left(\frac{\mathrm{d}}{\mathrm{d}t}\right)^{k-1} (1-t^2)^{k+(D-3)/2}\Bigg|_\theta^1 \\
&= -C\left(\frac{\mathrm{d}}{\mathrm{d}t}\right)^{k-1} (1-t^2)^{k+(D-3)/2}\Bigg|_{t=\theta} \qquad (k \geq 1)
\end{aligned}$$

When $k = 0$ we find that

$$\begin{aligned}
I_{0,0}(\theta) &= \int_\theta^1 (1-t^2)^{(D-3)/2}\mathrm{d}t \\
&= t\ {}_2F_1(1/2,(3-D)/2;3/2;t^2)\big|_\theta^1 \\
&= \frac{\sqrt{\pi}\,\Gamma((D-1)/2)}{2\Gamma(D/2)} - \theta\ {}_2F_1(1/2,(3-D)/2;3/2;\theta^2).
\end{aligned}$$

For $\alpha = 1$, we integrate by parts once and find that for $k \geq 2$,

$$
\begin{aligned}
I_{1,k}(\theta) &= C \int_\theta^1 t \left(\frac{\mathrm{d}}{\mathrm{d}t}\right)^k (1-t^2)^{k+(D-3)/2} \mathrm{d}t \\
&= C \left[ t \left(\frac{\mathrm{d}}{\mathrm{d}t}\right)^{k-1} (1-t^2)^{k+(D-3)/2} \Big|_\theta^1 - \int_\theta^1 \left(\frac{\mathrm{d}}{\mathrm{d}t}\right)^{k-1} (1-t^2)^{k+(D-3)/2} \mathrm{d}t \right] \\
&= C \left[ t \left(\frac{\mathrm{d}}{\mathrm{d}t}\right)^{k-1} (1-t^2)^{k+(D-3)/2} \Big|_\theta^1 - \left(\frac{\mathrm{d}}{\mathrm{d}t}\right)^{k-2} (1-t^2)^{k+(D-3)/2} \Big|_\theta^1 \right] \\
&= C \left[ \left(\frac{\mathrm{d}}{\mathrm{d}t}\right)^{k-2} (1-t^2)^{k+(D-3)/2} - t \left(\frac{\mathrm{d}}{\mathrm{d}t}\right)^{k-1} (1-t^2)^{k+(D-3)/2} \right] \Bigg|_{t=\theta} \qquad (k \geq 2)
\end{aligned}
$$

When $\alpha = 0$, we have a straightforward integral

$$
\begin{aligned}
I_{1,0}(\theta) &= \int_\theta^1 t(1-t^2)^{(D-3)/2} \mathrm{d}t \\
&= \frac{(1-\theta^2)^{(D-1)/2}}{(D-1)} = I_{0,1}(\theta).
\end{aligned}
$$

Finally, for $k = 1$, we obtain

$$
\begin{aligned}
I_{1,1}(\theta) &= \int_\theta^1 t^2(1-t^2)^{(D-3)/2} \mathrm{d}t \\
&= (t^3/3)\ {}_2F_1(3/2, (3-D)/2; 5/2; t^2)\big|_\theta^1 \\
&= \frac{\sqrt{\pi}\Gamma((D-1)/2)}{4\Gamma((D+2)/2)} - (\theta^3/3)\ {}_2F_1(3/2, (3-D)/2; 5/2; \theta^2)
\end{aligned}
$$

##### C.3.2 Properties of the eigenvalues of $K_{\text{shell}}$

The above show that for $k \geq 2$

$$
a_k = |\mathbb{S}^{D-2}|(I_{1,k}(\theta) - \theta I_{0,k}(\theta)) \propto \left(\frac{\mathrm{d}}{\mathrm{d}t}\right)^{k-2} (1-t^2)^{k+(D-3)/2} \Bigg|_{t=\theta}. \tag{50}
$$

Taking $\theta = -1$ leads to $a_k = 0$, since fewer derivatives than $k + (D-3)/2$ appear in (50), which reflects the fact that higher degree ultraspherical polynomials are orthogonal to a linear function. Furthermore, since $1 - t^2$ is an even function, the parity of $a_k$ as a function of $\theta$ matches the parity of $k$. However, $a_k$ appears squared in (48), so $\xi_k$ will always be an even function of $\theta$. This explains the parity symmetry of the eigenvalues with coding level for $k \geq 2$. Also, (50) for $\theta = 0$ gives $a_k = 0$ when $k$ is odd, as was shown by Bach (2017); Basri et al. (2019). This is because

$$\begin{aligned}
\left(\frac{\mathrm{d}}{\mathrm{d}t}\right)^p (1-t^2)^{p+\ell}\bigg|_{t=0} &= \left(\frac{\mathrm{d}}{\mathrm{d}t}\right)^p (1-t)^{p+l}(1+t)^{p+l}\bigg|_{t=0} \\
&= \sum_{j=0}^{p} \binom{p}{j} \left(\left(\frac{\mathrm{d}}{\mathrm{d}t}\right)^j (1-t)^{p+l}\right)\left(\left(\frac{\mathrm{d}}{\mathrm{d}t}\right)^{p-j} (1+t)^{p+l}\right)\bigg|_{t=0} \\
&= \sum_{j=0}^{p} \binom{p}{j} (-1)^j \left(\left(\frac{\mathrm{d}}{\mathrm{d}t}\right)^j (1+t)^{p+l}\right)\left(\left(\frac{\mathrm{d}}{\mathrm{d}t}\right)^{p-j} (1+t)^{p+l}\right)\bigg|_{t=0} \\
&= 0 \text{ if } p \text{ is odd,}
\end{aligned}$$

because the $j$ and $p-j$ terms have opposite parity and cancel.

We may also compute the tail asymptotics of these eigenvalues for large $k$. Let $p = k-2$ and $\ell = (D+1)/2$, so we want to evaluate

$$\begin{aligned}
\left(\frac{\mathrm{d}}{\mathrm{d}t}\right)^p (1-t^2)^{p+\ell} &= \frac{p!}{2\pi i}\oint \frac{(1-z^2)^{p+\ell}}{(z-t)^{p+1}}\mathrm{d}z \\
&= \frac{p!}{2\pi i}\oint e^{(p+o(p))F(z))}\mathrm{d}z
\end{aligned}$$

for large $p$ at $t = \theta \in (-1,1)$. The first line follows from Cauchy's intergral formula for a counterclockwise contour encircling $t$, and the second comes from defining

$$\begin{aligned}
F(z) &:= \log(1-z^2) - \log(z-t) \\
&\sim (1+\ell/p)\log(1-z^2) - (1+1/p)\log(z-t),
\end{aligned}$$

when $p$ is large and $\ell$ is constant. We will use the saddle point method (Butler, 2007) to evaluate the contour integral asymptotically, ignoring the $o(p)$ term in the exponent. Note that the only singularity in the original integrand occurs at $z = t$.

The function $F$ has derivatives

$$\begin{aligned}
F'(z) &= \frac{-2z}{1-z^2} - \frac{1}{z-t}, \\
F''(z) &= \frac{1}{(z-t)^2} - \frac{4z^2}{(1-z^2)^2} - \frac{2}{1-z^2}.
\end{aligned}$$

We find the saddle points by setting $F'(z) = 0$. This leads to a quadratic equation with two roots: $z_\pm = t \pm \sqrt{t^2 - 1} = \mathrm{sgn}(t)(|t| \pm i\sqrt{1-t^2})$. Since these are evaluated at $t = \theta$ with $|\theta| < 1$, both roots are complex, $|z_\pm| = 1$, and $F''(z_\pm) \neq 0$. Also, the saddle points avoid the singularity in the original integrand, so we can deform our contour to pass through these points and apply the standard approximation.

Applying the saddle point approximation, we obtain

$$
\begin{aligned}
\left(\frac{\mathrm{d}}{\mathrm{d}t}\right)^p (1-t^2)^{p+\ell} &\simeq \frac{p!}{2\pi i} \oint e^{pF(z)} \mathrm{d}z \\
&\simeq \frac{p!}{2\pi i} \sum_{z_0 \in \{z_+, z_-\}} e^{pF(z_0)} e^{i(\pi - \arg F''(z_0))/2} \left(\frac{2\pi}{p|F''(z_0)|}\right)^{1/2} \\
&\leq cp! \sum_{z_0 \in \{z_+, z_-\}} e^{pF(z_0)} p^{-1/2} \\
&= cp!\, p^{-1/2} \left( \left(\frac{1 - z_+^2}{z_+ - t}\right)^p + \left(\frac{1 - z_-^2}{z_- - t}\right)^p \right) \\
&= cp!\, p^{-1/2} \left((-2z_+)^p + (-2z_-)^p\right) \\
&\leq 2cp!\, p^{-1/2} (-2)^p
\end{aligned}
$$

for some $c$ which is constant in $p$ and depends on $D$. In the last step, we use that $z_+^p + z_-^p \leq 2$ since $z_\pm$ are conjugate pairs with magnitude 1.

Now recall the full equation for the coefficients:

$$
a_k = |\mathbb{S}^{D-2}|(-1/2)^k \frac{\Gamma((D-1)/2)}{\Gamma(k + (D-1)/2)} \left(\frac{\mathrm{d}}{\mathrm{d}t}\right)^p (1-t^2)^{p+\ell}.
$$

Plugging in the result from the saddle point approximation, substituting $p = k - 2$, and dropping all terms that are constant in $k$, we find that

$$
\begin{aligned}
a_k &\leq C'(-1/2)^k \frac{(k-2)!\, k^{-1/2}(-2)^k}{\Gamma(k + (D-1)/2)} \\
&= C' k^{-1/2} \frac{\Gamma(k-1)}{\Gamma(k + (D-1)/2)} \\
&= C' k^{-D/2-1},
\end{aligned}
$$

where $C'$ is a new constant. The rate of $k^{-D/2-1}$ is the same decay rate found by Bach (2017); Bietti and Bach (2021) using a different mathematical technique for $\theta = 0$. These decay rates are important for obtaining general worst-case bounds for kernel learning of general targets; Bach (2012) is an example.

#### C.4 Gaussian process targets

Taking our target function to be a GP on the unit sphere $f(\mathbf{x}) \sim \mathrm{GP}(0, C)$ with some covariance function $C : \mathbb{S}^{D-1} \times \mathbb{S}^{D-1} \to \mathbb{R}$, we can represent our target function by performing an eigendecomposition of the covariance operator $\mathcal{U}_C$. When $C$ itself is spherically symmetric and positive definite, this becomes

$$C(\mathbf{x}^\mu, \mathbf{x}^\nu) = \sum_{k=0}^{\infty} \rho_k \sum_{m=1}^{N(D,k)} Y_{km}(\mathbf{x}^\mu) Y_{km}(\mathbf{x}^\nu), \tag{51}$$

where $\rho_k > 0$ are the eigenvalues. Then a sample from the GP with this covariance function is a random series

$$f(\mathbf{x}) = \sum_{k=0}^{\infty} \sqrt{\rho_k} \sum_{m=1}^{N(D,k)} g_{km} Y_{km}(\mathbf{x}), \tag{52}$$

where $g_{km} \sim \mathcal{N}(0,1)$ by the Kosambi-Karhunen–Loève theorem (Kosambi, 1943). In other words, the coefficient of $Y_{km}$ in the series expansion of $f(\mathbf{x})$ is $c_{km} = \sqrt{\rho_k} g_{km}$.

We take the squared exponential covariance on the sphere

$$C(\mathbf{x}^\mu, \mathbf{x}^\nu) = \exp\left( \frac{-\|\mathbf{x}^\mu - \mathbf{x}^\nu\|^2}{2\gamma^2} \right) = \exp\left( \frac{t-1}{\gamma^2} \right), \tag{53}$$

for $t = \mathbf{x}^\mu \cdot \mathbf{x}^\nu$ and length scale $\gamma$.

#### C.5 Numerical details

All of our spherical harmonic expansions are truncated at frequency $N_k$. This is typically $N_k = 50$ for experiments in $D = 3$ dimensions. In higher dimensions, $N(D, k)$ grows very quickly in $k$, requiring truncation at a lower frequency.

To compute the kernel eigenvalues $\lambda_k$, we can either numerically integrate the Funk-Hecke formula (41) or compute the coefficients $a_k(\theta/r)$ semi-analytically, following (50), then integrate (48) with numerical quadrature and rescale by (34).

We use the Funk-Hecke formula (41) and numerical quadrature to find $\rho_k$. To compute the expected error using (54), we use $\mathbb{E}[c_\alpha^2] = \rho_k$. After generating a sample from the GP, we normalize the functions by dividing the labels and coefficients by their standard deviation. This ensures that the relative mean squared error (11) is equivalent to the mean squared error computed in the next section.

### D Calculation of generalization error

The generalization error of kernel ridge regression is derived in Canatar et al. (2021b); Simon et al. (2021); Gerace et al. (2020), which show that the mean squared error, in the absence of noise in the target, can be written as[2]

$$\mathbb{E}_{\mathbf{x}} (f(\mathbf{x}) - \hat{f}(\mathbf{x}))^2 = \sum_\alpha \beta_\alpha c_\alpha^2, \tag{54}$$

where $\beta_\alpha$ depend on $P$ and the kernel but not on the target, and $c_\alpha$ are the coefficients (37) of the target function in the basis $L^2(\sigma)$. Specifically,

$$\beta_\alpha = \left(\frac{1}{1-\chi}\right)\left(\frac{\kappa}{\lambda_\alpha P + \kappa}\right)^2, \tag{55}$$

where $\alpha$ indexes the kernel eigenfunctions and

$$\chi = \sum_\alpha \frac{\lambda_\alpha^2 P}{(\lambda_\alpha P + \kappa)^2}, \tag{56}$$

$$\kappa = \alpha_{\text{ridge}} + \sum_\alpha \frac{\lambda_\alpha \kappa}{\lambda_\alpha P + \kappa}, \tag{57}$$

with $\alpha_{\text{ridge}}$ the ridge parameter. Note that (57) is an implicit equation for $\kappa$, which we solve by numerical root-finding.

Thus,

$$\mathbb{E}_{\mathbf{x}} (f(\mathbf{x}) - \hat{f}(\mathbf{x}))^2 = C_1 \sum_\alpha \left(\frac{c_\alpha}{\lambda_\alpha + C_2}\right)^2, \tag{58}$$

with $C_1 = \left(\frac{1}{1-\chi}\right)\frac{\kappa^2}{P^2}$ and $C_2 = \frac{\kappa}{P}$.

[2]The exact form of this expression differs from that given in Canatar et al. (2021b) due to differences in the conventions we take for our basis expansions.

### E Dense-sparse networks

To compare with more realistic networks, we break the simplifying assumption that $\mathbf{J}^{\text{eff}}$ is densely connected and instead consider sparse connections between the input and expansion layer. Consider a random matrix $\mathbf{J}^{\text{eff}} = \mathbf{J}\mathbf{A}$, where $\mathbf{A} \in \mathbb{R}^{N \times D}$ and $\mathbf{J} \in \mathbb{R}^{M \times N}$, with $N > D$ and $M > N$. The entries of $\mathbf{A}$ are i.i.d. Gaussian, i.e. $A_{ij} \sim \mathcal{N}(0, 1/D)$. In contrast, $\mathbf{J}$ is a sparse matrix with *exactly $K$ nonzero entries per row*, and nonzero entries equal to $1/\sqrt{K}$. With these scaling choices, the elements of $\mathbf{J}^{\text{eff}}$ are of order $1/\sqrt{D}$, which is appropriate when the input features $x_i$ are order 1. This is in contrast to the rest of this paper, where we considered features of order $1/\sqrt{D}$ and therefore assumed order 1 weights. The current scaling allows us to study the properties of $\mathbf{J}^{\text{eff}}$ for different values of $D$, $N$ and $K$.

First, we examine properties of $\mathbf{J}^{\text{eff}} = \mathbf{J}\mathbf{A}$ under these assumptions. Recall that the rows of $\mathbf{J}^{\text{eff}}$ are the weights of each hidden layer neuron. First, since $\mathbf{A}$ is Gaussian, any given row $\mathbf{J}_i^{\text{eff}} \in \mathbb{R}^D$ is marginally Gaussian and distributed identically to any other row. But the rows are *not* independent, since they are all linear combinations of the rows of $\mathbf{A}$. Thus, the kernel limit of an infinitely large dense-sparse network is equal to that of a fully dense network, but convergence to that kernel behaves differently and requires taking a limit of both $N, M \to \infty$. In this section, we study how finite $N$ introduces extra correlations among the rows of $\mathbf{J}^{\text{eff}}$ compared to dense networks.

The distribution of $\mathbf{J}^{\text{eff}}$ is spherically symmetric in the sense that $\mathbf{J}^{\text{eff}}$ and $\mathbf{J}^{\text{eff}}\mathbf{Q}$ have the same distribution for any rotation matrix $\mathbf{Q} \in \mathbb{R}^{D \times D}$. In contrast, a densely connected network with weights $\mathbf{G}$ drawn i.i.d. as $G_{ij} \sim \mathcal{N}(0, 1/D)$ will of course have independent rows and also be spherically symmetric. The spherical Gaussian is the only vector random variable which is spherically symmetric with independent entries (Nash and Klamkin, 1976). Furthermore, each row of $\mathbf{G}$ may be rotated by a *different* orthogonal matrix and the resulting random variable would still have the same distribution.

With these symmetry considerations in mind, the statistics of the rows of $\mathbf{J}^{\text{eff}}$ can be described by their multi-point correlations. The simplest of these is the two-point correlation, which in the case of spherical symmetry is captured by the overlaps:

$$\nu_{ij} := \sum_{k=1}^{D} (\mathbf{J}^{\text{eff}})_{ik} (\mathbf{J}^{\text{eff}})_{jk} = \sum_{k=1}^{D} \sum_{m,n=1}^{N} J_{in} A_{nk} J_{jm} A_{mk} \ . \tag{59}$$

The overlap $\nu_{ij}$ is doubly stochastic: one source of stochasticity are the elements of $\mathbf{A}$, and the second one is the random sampling of nonzero elements of $\mathbf{J}$. Ideally, we are interested in studying the statistics of $\nu_{ij}$ when varying $i$ and $j$, i.e. when $\mathbf{J}$ varies (since the rows of $\mathbf{J}$ are sampled independently from each other). However, this will leave us with the quenched disorder given by the specific realization of $\mathbf{A}$. To obtain a more general and interpretable result, we want to compute the probability distribution

$$P_{\mathbf{A},\mathbf{J}}(\nu_{ij}) = \mathop{\mathbb{E}}_{\mathbf{A}}\left[\mathop{\mathbb{E}}_{\mathbf{J}}\left[\delta\left(\nu_{ij} - \sum_{k=1}^{D}\sum_{m,n=1}^{N} J_{in}J_{jm}A_{nk}A_{mk}\right)\right]\right] \quad . \tag{60}$$

Notice that the order in which we perform the averaging is irrelevant.

#### E.1 Computation of the moment-generating function

Instead of computing directly the probability distribution in Eq. 60, we compute the moment-generating function

$$Z(\mu) := \mathop{\mathbb{E}}_{\mathbf{A}}\left[\mathop{\mathbb{E}}_{\mathbf{J}}\left[\exp\left(\mu\nu_{ij}\right)\right]\right] \quad , \tag{61}$$

which fully characterizes the probability distribution of $\nu_{ij}$. We indicate the set of indices in which the $i$-th row of $\mathbf{J}$ takes nonzero values by $S^i = \{S^i_1, S^i_2, \ldots, S^i_K\}$ such that $J_{iS^i_l} \neq 0$, $\forall l = 1, \ldots, K$, and analogously for the $j$-th row. We also indicate the intersection $S^{ij} = S^i \cap S^j$, i.e. the set of indices in which *both* the $i$-th and the $j$-th rows are nonzero. $S^{ij}$ has size $0 \leq |S^{ij}| \leq K$. Notice that setting $i = j$ causes $|S^{ij}| = K$ deterministically. With this definitions, the overlap can be written as

$$\nu_{ij} = \sum_{k=1}^{D}\sum_{m\in S^i}\sum_{n\in S^j} J_{in}J_{jm}A_{nk}A_{mk} = \frac{1}{K}\sum_{k=1}^{D}\sum_{m\in S^i}\sum_{n\in S^j} A_{nk}A_{mk} \tag{62}$$

We start by perform swapping the averaging order in Eq. 61 and averaging over $\mathbf{A}$.

$$\begin{aligned}
Z(\mu) =& \mathop{\mathbb{E}}_{\mathbf{J}}\left[\int\left(\prod_{m=1}^{N}\prod_{l=1}^{D}\mathcal{D}A_{ml}\right)\exp\left(\frac{\mu}{K}\sum_{k=1}^{D}\sum_{m\in S^i}\sum_{n\in S^j}A_{nk}A_{mk}\right)\right] \\
=& \mathop{\mathbb{E}}_{\mathbf{J}}\left[\int\left(\prod_{m\in S^i\cup S^j}\prod_{l=1}^{D}\mathcal{D}A_{ml}\right)\exp\left(\frac{\mu}{K}\sum_{k=1}^{D}\sum_{m\in S^i}\sum_{n\in S^j}A_{nk}A_{mk}\right)\right] \\
=& \mathop{\mathbb{E}}_{\mathbf{J}}\left[\prod_{k=1}^{D}\int\left(\prod_{m\in S^i\cup S^j}\mathcal{D}A_{mk}\right)\exp\left(\frac{\mu}{K}\sum_{m\in S^i}\sum_{n\in S^j}A_{nk}A_{mk}\right)\right] \quad ,
\end{aligned}$$

where in the first equality we marginalized over all the elements of $\mathbf{A}$ which do not enter the definition of $\nu_{ij}$, i.e. we went from having to integrate over $N \times D$ variables to only $|S^i \cup S^j| \times D = (2K - |S^{ij}|) \times D$ variables. In the second equality we factorized the columns of $\mathbf{A}$.

We now explicitly compute integral for a fixed value of $k$, by reducing it to a Gaussian integral:

$$\begin{aligned}
&\int \left( \prod_{m \in S^i \cup S^j} \mathcal{D}A_{mk} \right) \exp\left( \frac{\mu}{K} \sum_{m \in S^i} \sum_{n \in S^j} A_{nk} A_{mk} \right) \\
&= \int \left( \prod_{m \in S^i \cup S^j} dA_{mk} \right) (2\pi D)^{-\frac{|S^i \cup S^j|}{2}} \exp\left( \frac{\mu}{K} \sum_{m \in S^i} \sum_{n \in S^j} A_{nk} A_{mk} - \frac{D}{2} \sum_{r \in S^i \cup S^j} A_{rk}^2 \right) \\
&= \int \left( \prod_{m \in S^i \cup S^j} dA_{mk} \right) (2\pi D)^{-\frac{|S^i \cup S^j|}{2}} \exp\left( -\frac{D}{2} \sum_{r \in S^i \cup S^j} A_{rk} P_{rs} A_{sk} \right) \\
&= \det(\mathbf{P})^{-1/2} \quad ,
\end{aligned}$$

where $\mathbf{P} \in \mathbb{R}^{|S^i \cup S^j| \times |S^i \cup S^j|}$, which has a 3-by-3 block structure and can be written as

$$\mathbf{P} = \begin{pmatrix} \mathbf{I}_{K-|S^{ij}|} & -\frac{\mu}{KD}\mathbf{1}_{|S^{ij}|} & -\frac{\mu}{KD}\mathbf{1}_{K-|S^{ij}|} \\ -\frac{\mu}{KD}\mathbf{1}_{|S^{ij}| \times (K-|S^{ij}|)} & \mathbf{I}_{|S^{ij}|} - 2\frac{\mu}{KD}\mathbf{1}_{|S^{ij}|} & -\frac{\mu}{KD}\mathbf{1}_{|S^{ij}| \times (K-|S^{ij}|)} \\ -\frac{\mu}{KD}\mathbf{1}_{K-|S^{ij}|} & -\frac{\mu}{KD}\mathbf{1}_{|S^{ij}| \times (K-|S^{ij}|)} & \mathbf{I}_{K-|S^{ij}|} \end{pmatrix} \quad , \tag{63}$$

where $I_n$ is the $n$-by-$n$ identity matrix and $\mathbf{1}_{n \times m}$ is the $n$-by-$m$ matrix of all ones (if $m$ is omitted, then it is $n$-by-$n$). Due to the block structure, the determinant of the matrix above is identical to the determinant of a 3-by-3 matrix

$$\begin{aligned}
\det(\mathbf{P}) &= \det \begin{pmatrix} 1 & -\frac{\mu}{KD}|S^{ij}| & -\frac{\mu}{KD}(K - |S^{ij}|) \\ -\frac{\mu}{KD}(K - |S^{ij}|) & 1 - 2\frac{\mu}{KD}|S^{ij}| & -\frac{\mu}{KD}(K - |S^{ij}|) \\ -\frac{\mu}{KD}(K - |S^{ij}|) & -\frac{\mu}{KD}|S^{ij}| & 1 \end{pmatrix} \\
&= \frac{K^2D^2 - K^2\mu^2 + |S^{ij}|^2\mu^2 - 2DK|S^{ij}|\mu}{K^2D^2} \quad .
\end{aligned} \tag{64}$$

By plugging this result into the expression for the moment-generating function, we have that

$$Z(\mu) = \mathop{\mathbb{E}}_{\mathbf{J}} \left[ \left( \frac{K^2D^2 - K^2\mu^2 + |S^{ij}|^2\mu^2 - 2DK|S^{ij}|\mu}{K^2D^2} \right)^{-D/2} \right] \quad . \tag{65}$$

This expression is our core result, and needs to be averaged over $\mathbf{J}$. This average can be written explicitly by noticing that, when $i \neq j$, $|S^{ij}|$ is a random variable that follows a hypergeometric distribution in which the number of draws is equal to number of success state and is equal to $K$. By using the explicit expression of the probability mass function of a hypergeometric distribution, we have that

$$Z(\mu) = \sum_{s=0}^{K} \frac{\binom{K}{s}\binom{N-K}{K-s}}{\binom{N}{K}} \left(\frac{K^2D^2 - K^2\mu^2 + s^2\mu^2 - 2DKs\mu}{K^2D^2}\right)^{-D/2} \quad . \tag{66}$$

Notice that the term $s = 0$ yields the same moment-generating function (up to a factor) as for a fully-connected $\mathbf{J}^{\text{eff}}$ with Gaussian i.i.d. entries with variance $1/D$. In contrast, when $i = j$ we obtain

$$Z_{i=j}(\mu) = \left(1 - \frac{2\mu}{D}\right)^{-D/2} \quad . \tag{67}$$

#### E.2 Computation of the moments of $\nu_{ij}$

In this section, we assume that $i \neq j$ and use the moment-generating function to compute the moments of $\nu_{ij}$. The non-central moments of the overlap are easily obtained from the moment-generating function as

$$\mathbb{E}\left[\nu_{ij}^q\right] = \frac{d^q}{d\mu^q} Z(\mu)|_{\mu=0} \quad , \tag{68}$$

which can be computed in a symbolic manipulation tool.

We now explicitly compute the first two moments of $\nu_{ij}$.

$$\mathbb{E}\left[\nu_{ij}\right] = \frac{d}{d\mu} Z(\mu)|_{\mu=0} = \frac{1}{K} \mathop{\mathbb{E}}_{\mathbf{J}}\left[|S^{ij}|\right] = \frac{K}{N} \quad , \tag{69}$$

where we used the fact that the mean of $s \sim \text{Hypergeom}(N, K, K)$ is given by $\mathbb{E}[s] = \frac{K^2}{N}$. For the second moment, we have

$$\begin{aligned}\mathbb{E}\left[\nu_{ij}^2\right] &= \mathop{\mathbb{E}}_{\mathbf{J}}\left[\frac{K^2 + |S^{ij}|^2(1+D)}{K^2D}\right] \\ &= \frac{1}{D} + \frac{D+1}{D}\left(\frac{(N-K)^2}{N^2(N-1)} + \frac{K^2}{N^2}\right)\end{aligned}$$

while to compute the variance we use the law of total variance

$$
\begin{aligned}
\mathrm{Var}(\nu_{ij}) &= \mathop{\mathbb{E}}_{s}[\mathrm{Var}(\nu_{ij}|s)] + \mathrm{Var}(\mathbb{E}\left[\nu_{ij}|s\right]) \\
&= \frac{1}{D} + \frac{D+1}{K^2 D}\,\mathbb{E}[s^2] - \frac{1}{K^2}\,\mathbb{E}[s^2] + \mathrm{Var}\left(\frac{1}{K}s\right) \\
&= \frac{1}{D} + \frac{D+1}{D}\left(\frac{(N-K)^2}{N^2(N-1)} + \frac{K^2}{N^2}\right) - \frac{K^2}{N^2} \\
&= \frac{1}{D} + \frac{D+1}{D}\left(\frac{(N-K)^2 + (N-1)K^2 - (N-1)K^2\frac{D}{D+1}}{N^2(N-1)}\right) \\
&= \frac{1}{D} + \frac{D+1}{D}\frac{1}{N-1} + \frac{1}{D}\frac{K^2 - 2(D+1)K}{N(N-1)} + \frac{K^2}{N^2(N-1)} \quad .
\end{aligned}
$$

As $N \to \infty$ we have that $\mathrm{Var}(\nu_{ij}) \sim \frac{1}{D}$, which is the same variance of the overlap for a fully-connected $\mathbf{J}^{\text{eff}}$ with Gaussian i.i.d. entries. This is expected since when $N$ is large, the probability of $i$ and $j$ having common afferents goes to zero.

#### E.3 Comparison to clustered embedding

Instead of distributed embedding, i.e. $\mathbf{A}$ being a Gaussian matrix, here we consider a clustered embedding by setting

$$
\mathbf{A} = \mathbf{I}_D \otimes \mathbf{1}_{N/D} \quad , \tag{70}
$$

i.e. the Kronecker product of the $D$-dimensional identity matrix and a vector of all ones and length $N/D$. This means that we can separate the input layer of $N$ neurons in $D$ non overlapping subsets $B_n = \{\frac{N}{D}(n-1)+1, \frac{N}{D}(n-1)+2, \ldots, \frac{N}{D}n\}$, each of size $N/D$, and we can write

$$
A_{mn} = \begin{cases} 1 & \text{if } m \in B_n \\ 0 & \text{otherwise} \end{cases} \quad . \tag{71}
$$

In this case the overlap is given by

$$
\nu_{ij} = \frac{1}{K}\sum_{l=1}^{D}\sum_{m\in S^i}\sum_{n\in S^j} 1\left[m \in B_l\right] 1\left[n \in B_l\right] \quad , \tag{72}
$$

where $1[\cdot]$ is the indicator function, i.e. it is one if the argument is true and zero if the argument is false. We indicate by $K_l^i$ the number of elements of $S^i$ which belongs to group

$l$, i.e. $K_l^i = \sum_{m \in S^i} 1[m \in B_l]$. The overlap can then be written as

$$\nu_{ij} = \frac{1}{K} \sum_{l=1}^{D} K_l^i K_l^j \quad . \tag{73}$$

The vector $\mathbf{K}^i = (K_1^i, \ldots, K_D^i)$ follows a multivariate hypergeometric distribution with $D$ classes, $K$ draws, a population of size $N$, and number of successes for each class equal to $N/D$. Notice that $\mathbf{K}^i$ and $\mathbf{K}^j$ are independent from each other since each neuron samples its pre-synaptic partners independently. We can now compute explicitly the mean of $\nu_{ij}$ using the fact that $\mathbb{E}\left[K_l^i\right] = \frac{K}{D}$

$$\mathbb{E}\left[\nu_{ij}\right] = \frac{1}{K} \sum_{l=1}^{D} \mathbb{E}\left[K_L^i\right]^2 = \frac{K}{D} \quad . \tag{74}$$

Similarly, we can write the second moment of $\nu_{ij}$ as

$$\mathbb{E}\left[\nu_{ij}^2\right] = \frac{1}{K^2} \left( \sum_{l=1}^{D} \mathbb{E}\left[(K_l^i)^2\right]^2 + \sum_{l \neq l'} \mathbb{E}\left[K_l^i K_{l'}^i\right]^2 \right) \quad . \tag{75}$$

Once again, we can use known result for variance and covariance of multivariate hypergeometric variables to simplify the above expression. Indeed, we can write

$$\mathbb{E}\left[(K_l^i)^2\right] = K \frac{N-K}{N-1} \frac{D-1}{D^2} + \frac{K^2}{D^2} \tag{76}$$

$$\mathbb{E}\left[K_l^i K_{l'}^i\right] = -K \frac{N-K}{N-1} \left(\frac{1}{D}\right)^2 + \frac{K^2}{D^2} \tag{77}$$

from which we obtain the final expression for the second moment

$$\mathbb{E}\left[\nu_{ij}^2\right] = \frac{K^2}{D^2} + \frac{1}{D}\left(1 - \frac{1}{D}\right)\left(\frac{N-K}{N-1}\right)^2 \quad . \tag{78}$$
